# kinfitr: Reproducible PET Pharmacokinetic Modelling in R

**DOI:** 10.1101/755751

**Authors:** Granville J. Matheson

## Abstract

Quantification of Positron Emission Tomography (PET) data is performed using pharmacokinetic models. There exist many models for describing this data, each of which may describe the data better or worse depending on the specific application, and there are both theoretical, practical and empirical reasons to select any one model over another. As such, effective PET modelling requires a high degree of flexibility, while effective communication of all steps taken through scientific publications is not always feasible. Reproducible research practices address these concerns, in that researchers share analysis code, and data if possible, such that all steps are recorded, allowing an independent researcher to reproduce the results and assess their veracity. In this article, I present *kinfitr*: a software package for performing kinetic modelling using the open-source R language, in a reproducible manner. The R community has a strong culture of reproducible research, and the language consists of numerous tools which allow both effective and easy sharing and communication of analysis code. The package is written in such a way as to allow the analyst the freedom to use and rapidly exchange between approaches, and to assess goodness of fit, with 14 different kinetic models currently implemented using a consistent syntax, as well as tools for working with the data. By providing open-source tools for kinetic modelling, including documentation and examples, it is hoped that this will extend access to methodology for research groups lacking software engineering expertise, as well as simplify and thereby encourage transparent and reproducible reporting.

## INTRODUCTION

Positron emission tomography (PET) is an *in vivo* neuroimaging method with high biochemical sensitivity and specificity: it is an essential tool for the study of the neurochemical pathophysiology of mental and neurological disease as well as for evaluating pharmacological treatments. This method allows for accurate quantification of picomolar concentrations, thereby allowing insights which are not possible using any other in vivo imaging modality. PET is, however, prohibitively expensive, often costing in excess of USD 10 000 per measurement, and additionally involves exposure of participants to harmful radioactivity. For this reason, accurate quantification is imperative in order to maximise the scientific value of each measurement, as well as to minimise the number of participants who must be exposed to radiation in order to answer the scientific question at hand.

Receptor PET quantification involves measuring the radioactivity in the tissue of interest following the injection of a radiolabelleled ligand into the blood. For fully quantitative PET imaging, radioactivity concentrations are measured over time, giving rise to a time activity curve (TAC). By examining the dynamics of radioactivity concentrations as the ligand enters and exits the tissue, the researcher is able to fit a model with which she can obtain information about the pharmacokinetics and binding of the ligand, and thereby assess the concentration of the relevant protein. The most common use of PET modelling involves the quantification of a static quantity of interest, which is estimated using a model fitted to the entire dynamic time-course of measurements in any given region. This is in contrast with fMRI, for which each 3D image within the time series represents the quantity of interest. The model used to perform PET quantification is therefore of tremendous importance for the later statistical comparison using the estimated quantity of interest.

There exist numerous different kinetic models for performing this quantification, which differ in a variety of important ways. Firstly, they differ in their specificity for the target binding, e.g. quantifying only the specific binding itself, as compared to quantifying the total binding, including non-specific binding (Innis et al. 2007). Secondly, they differ in their level of detail in their output, e.g. estimating only the estimate of the binding, compared to estimating all of the rate constants underlying that estimate. Thirdly, they differ in their relative degree of bias and variance, and hence their sensitivity to noise, i.e. how much they over- or under-fit the data). Fourthly, they differ in their assumptions about the behaviour of the radiotracer in the tissue, e.g. irreversible vs reversible binding, or the compartmental structure of the binding (Figure 1). These assumptions are usually only partially met in any given application, and care must be taken to ensure that the degree to which assumptions are not met does not bias the estimates in important ways (Salinas, Searle, and Gunn 2014). These differences are further complicated by the fact that the performance of different models may vary based on the properties of each specific tracer: a certain model may be more effective for certain cases than others, all else being equal.

**Figure 1.**
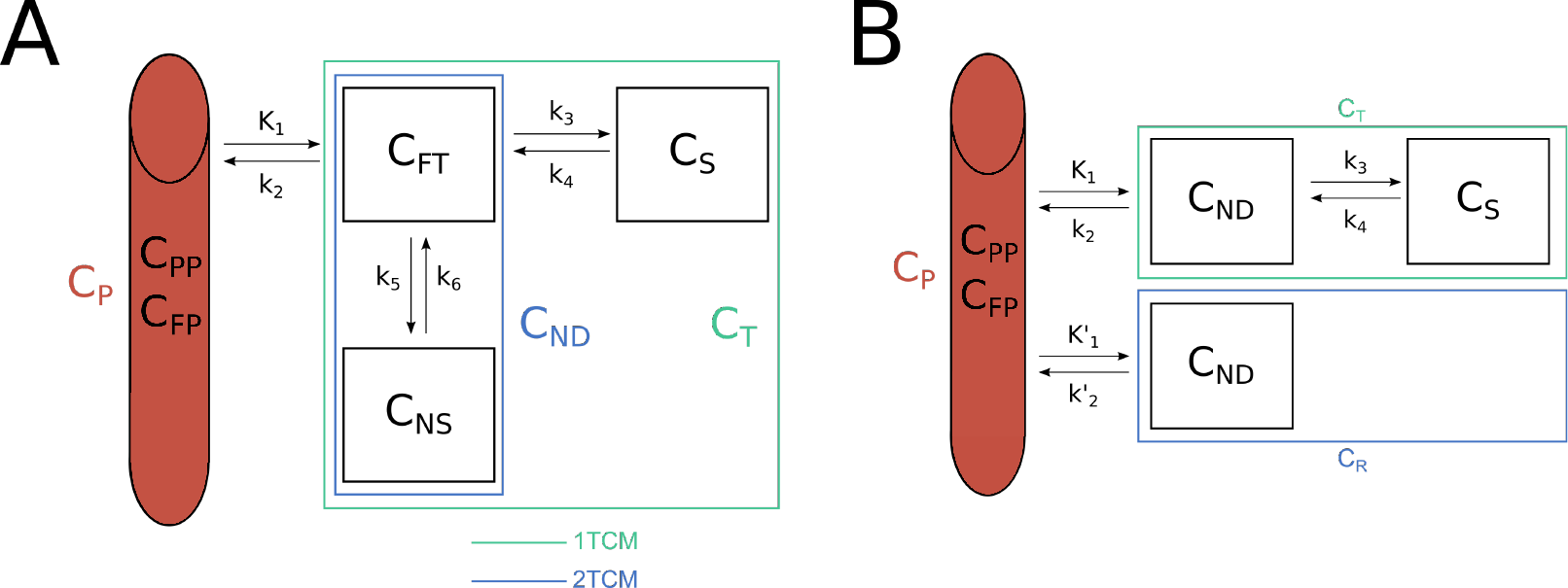
Compartmental models are the basis of PET kinetic modelling. For both panels, C represents the radioactivity concentrations within each compartment. The red cylinder on the left of each panel represents the arterial blood, containing plasma (P). Within the plasma, the radiotracer is either free (FP), or bound to plasma proteins (PP). The black boxes represent the compartments. TCM refers to Tissue Compartment Model. A. The three tissue compartment model is the basis for the two- and one-tissue compartment models: transfer between certain compartments are assumed to be sufficiently rapid that they can be considered as single compartments for the two- and one-tissue compartment models (coloured boxes). The compartments include FT free tracer, NS non-specifically bound, S specifically bound, T total, and ND non-displaceable. B. Reference region models consider the total concentration of radiotracer in the target T and in the reference region R, and assume that the non-displaceable concentration is equal in both regions, and that the specific binding in the reference region is equal to 0.

For the modeller, there is no silver bullet. Rather, the model used to estimate the quantity of interest should be selected based on the radiotracer, as well as the research question and properties of the data set itself. This is further complicated by the myriad other analytical decisions which must be made prior to modelling, such as statistical weighting schemes, the application of partial volume effect correction, or the use of numerous ways that the blood data can be modelled too (the blood, blood-to-plasma ratio and parent fraction curves can all be modelled to derive improved estimates of the arterial input function curve, which can itself also be modelled). As such, effective PET modelling requires a high degree of flexibility, and the ability to rapidly exchange between different models. Effective communication of all steps taken through scientific publications, however, is not always feasible due to the large number of small decisions and results which are sometimes required to reach a decision about how best the data should be modelled. This complicates replication efforts and thereby retards scientific progress. This problem is by no means restricted to PET modelling, and is a property of the increasing complexity of computational analysis of scientific data more generally.

This general issue has led to calls among the broader scientific community for computational reproducibility, or more broadly *reproducible research* (RR), as a minimum standard for assessment of scientific claims, i.e. that researchers share analysis code and, if possible, data. This ensures not only that all steps are recorded, but this also allows an independent researcher to reproduce the results and assess their veracity, as well as their sensitivity to various decisions taken during analysis. RR practices further accelerate scientific progress, as novel methods can be readily validated, applied and extended by other researchers using the shared code.

In this paper, I present *kinfitr*: a software package for performing PET kinetic modelling of TACs using the R language. This tool both provides flexibility for effective modelling, while at the same time being written in such a way as to promote transparency of this process. Further, by using the R language, all code is open-source, and reproducible reporting is made easy by the extensive ecosystem of tools for this purpose for the R language.

## DESCRIPTION

The *kinfitr* package contains a host of tools for processing and modelling of PET TAC data, i.e. after the raw image data has been transformed into vectors of radioactivity concentrations. The code is available at https://github.com/mathesong/kinfitr.

### Model Functions

There currently exist 14 models for TAC modelling (Gunn, Gunn, and Cunningham 2001; Rizzo et al. 2014; Logan et al. 1990, 1996; Ichise et al. 2002, 2003; Turkheimer et al. 2003; Patlak, Blasberg, and Fenstermacher 1983; Patlak and Blasberg 1985; Todd Ogden, Zanderigo, and Parsey 2015; Lammertsma and Hume 1996; Tomasi et al. 2008), including models which quantify binding relative to arterial plasma input, reference regions, or semi-quantitative methods by which binding is quantified relative to the injected dose of radioactivity and the mass of the participant. The models included and their descriptions are included in Table 1. All models have associated plotting routines, which include representation of weights, and the corresponding reference curve. Most models also produce standard errors of estimates, estimated using the delta method when the parameters are not directly fitted.

**Table 1.** Kinetic models included in the package for modelling of time activity curves

| **Model** | **(Ir)reversible** | **Fit Method** | **t\*** | **Input** | **Fit Parameters** | **Reference** |
| --- | --- | --- | --- | --- | --- | --- |
| One Tissue Compartment Model | Either | NLS |  | AIF | 2-4 |  |
| Two Tissue Compartment Model | Either | NLS |  | AIF | 4-6 |  |
| Two Tissue Compartment Model with Irreversible Trapping | Either | NLS |  | AIF | 5-7 | Rizzo et al., 2014 |
| Logan Plot | Reversible | OLS | Required | AIF | 1 | Logan et al., (1990) |
| Ichise Multilinear Analysis 1 | Reversible | OLS | Required | AIF | 1 | Ichise et al., (2002) |
| Ichise Multilinear Analysis 2 | Reversible | OLS | Required | AIF | 1 | Ichise et al., (2002) |
| Multilinear Logan Plot | Reversible | OLS | Required | AIF | 1 | Turkheimer et al., (2003) |
| Patlak Plot | Irreversible | OLS | Required | AIF | 1 | Patlak et al., (1983) |
| Simultaneous Estimation of Non-Displaceable Binding | Reversible | NLS + GridSearch |  | AIF | 1 | Ogden et al., (2015) |
| Simplified Reference Tissue Model | Reversible | NLS |  | Reference Tissue | 3 | Lammertsma & Hume, (1996) |
| Simplified Reference Tissue Model with Blood Volumes | Reversible | NLS |  | Reference Tissue | 4-5 | Tomasi et al., (2008) |
| Non-Invasive Logan Plot | Reversible | OLS | Required | Reference Tissue | 1 | Logan et al., (1996) |
| Ichise's Multilinear Reference Tissue Model | Reversible | OLS | Optional | Reference Tissue | 2 | Ichise et al., (2003) |
| Ichise's Multilinear Reference Tissue Model 2 | Reversible | OLS | Optional | Reference Tissue | 1 | Ichise et al., (2003) |
| Non-Invasive Multilinear Logan Plot | Reversible | OLS | Required | Reference Tissue | 1 | Turkheimer et al., (2003) |
| Patlak Reference Tissue Model | Irreversible | OLS | Required | Reference Tissue | 1 | Patlak et al., (1985) |

### Model Fitting

In addition to the standard implementations of all models (ordinary least squares for linear models, and Levenberg-Marquardt algorithm for nonlinear models), all those which are fitted using nonlinear least squares (NLS) have an additional option to be fit multiple times using different starting parameters: this is useful when there are local minima within parameter space. Using the *nls.multstart* package (Padfield and Matheson 2018), starting parameters can either be sampled either from a uniform distribution, or from a grid, across parameter space to ensure that the best-fitting parameters are identified.

For most linear models, there is a t* value which must be supplied. This value represents the point at which linearity is reached. For all models for which a t* value can be provided, there are included functions for selecting which t* value is most appropriate. As an intentional design decision, the selection of the t* is not automated: rather, the user is presented with R^2^ values, percentage variance and changes in binding estimates for each potential t* value such that an appropriate value can be selected from examining plots for several individuals to determine a t* value for a sample.

### Blood preprocessing

The *kinfitr* package contains a set of tools for preprocessing of blood data. Blood data can be read into *kinfitr* directly and automatically from PET BIDS JSON files as *blooddata* objects. This contains the raw data for the arterial blood, arterial plasma, blood-to-plasma ratio, parent fraction, and arterial input function, as well as the models used to interpolate this data. These objects also contain the model which will be used to interpolate this data. By default, the interpolation method is defined as piecewise linear interpolation, but all curves can also be interpolated using nonlinear models. All of the data points, interpolated curves and their combination to produce the final arterial input function can be visualised using a *plot* command.

There are numerous nonlinear models available for the modelling blood curves, and a selection of these models are included in the *kinfitr* package. For modelling of the parent fraction, the Hill, power, exponential and cumulative inverse gamma functions models are included (Tonietto et al. 2016), as well as the modification of the Hill function by Guo et al. (Guo et al. 2013). For modelling of blood or arterial input function curves, the tri-exponential model (linear increase, followed by a tri-exponential decay), as well as a spline model are available. Due to the large number, and *ad hoc* nature of many models for modelling of blood curves, it is also possible to create new models which can be incorportated into the *blooddata* object, or even to use the predictions from another external model (e.g. from another piece of software).

Once the user is satisfied with the fits to the blood data, an *input* object can be created: this consists of an interpolation of the curves into a common time series which can be used for kinetic modelling. Additionally, if the user only has access to blood data which is already preprocessed, they can also bypass the *blooddata* object, and directly interpolate the data into an *input* object.

### Included Data and Vignettes

The *kinfitr* package contains two data sets in order to allow for the inclusion of examples with most functions for demonstrating their uses. The first dataset, called *pbr28*, consists of TACs and blood data from several individuals measured using [^11^C]PBR28 from Matheson et al. (2017). This data includes 10 individuals, each measured twice in a test-retest study protocol, and includes TACs from six regions as well as blood data. Blood data is provided both in processed form, as was used in Matheson et al. (2017), but also in raw BIDS format. The second dataset, called *simref*, consists of a simulated dataset of 20 individuals which can be modelled using a reference region approach.

The package will also contain several vignettes which are currently under development, demonstrating how the functions of the package can be used. This allows new users to quickly learn how to use the package, and to allow them to get started by recycling, and reverse-engineering already-written code, rather than beginning from scratch. The included vignettes will be as follows:

1. Reference Tissue Models
2. Arterial Input Models
3. Choosing a Suitable t* in Linear Models
4. Pre-processing and Modelling of Blood Data

#### Other Fuctions

The package also contains a number of helper functions which can be used for the kinds of calculations and processesing steps which must often be performed to accompany TAC modelling. The package includes a unit conversion function, for translating between any standard units of radioactivity to any other, as well as for applying, and reversing, decay correction. For blood data collected using automated blood sampling systems, the package offers dispersion correction. The package additionally methods for estimating weights of TACs.

For models involving arterial input, the TAC data and arterial input data must be matched in time. All of the “tissue compartment models” allow for additional fitting of the time delay between these curves, as well as the blood volume fraction as additional parameters.

### Inputs and Syntax

All tools within the *kinfitr* package have been designed to function with numeric vectors to as great an extent as possible, as opposed to highly structured lists. This is an intentional design decision in order to allow most functions within the package not to be reliant on having created specific data structures in previous steps. Instead, users should be able to make use of functions from the package at whichever stage of a given processing pipeline. Notable exceptions are those of *blooddata* and *input* objects, which are created to make it easier to deal with blood data arising from a multitude of different sources, with different time sequences.

Modelling functions have been written in such a way as to be as consistent with one another as possible. Input arguments are the same between functions, and can be copied between the different models. This allows users to quickly switch between different models with mostly, if not completely, the same input arguments, thereby allowing a high degree of flexibility.

### Output

Fit objects are created in order to be as extensive as possible. The specific model fits themselves are included in the fit object, allowing them to be probed using methods such as AIC (Akaike Information Criterion), BIC (Bayesian Information Criterion), vcov (return the variance-covariance matrix), afforded by the stats package within the base R language. Additionally, *input*, weights and TAC data are included after their time shifting to correspond with one another, as well as predicted values including which points are to be considered before and after the t* point for those models which contain a t*. While this is memory-intensive, this step allows users to rapidly identify the causes of a poor fit.

### The R Programming Language and Reproducibility

One of the primary benefits of *kinfitr* is its being situated within the R programming language. R is an open-source programming (R Core Team 2019) language designed by and for statisticians, and one of the dominant languages used in Data Science. As such, it is excellent for performing data cleaning and rearrangement before modelling, as well as for later statistical analysis and data visualisation. Further, it makes it especially easy to implement additional methods which might be required to supplement the tools available within *kinfitr* as a result of CRAN (the Comprehensive R Archive Network): the central package repository for the R language, containing nearly 15 000 additional open-source packages.

Another key advantage of *kinfitr* being situated within the R programming language is access to the extensive collection of tools for ensuring reproducibility. Rmarkdown (Xie 2017) allows for code, code output (including plots) and text to be written side-by-side such that a new document can be generated when the data changes. This is not limited to analysis reports: entire scientific articles (such as this one) can be written within R (Allaire et al. 2019), where the plots and tables and even the values within the text can be automatically updated when the document is re-compiled. While this may takes more time when first creating the report, the time savings in the long run can be dramatic, as all changes based on alterations to processing or the underlying data are automatically incorporated. Furthermore, this is an effective strategy to minimise errors when transferring figures between software, and, by being scripted, allows for rapid identification and diagnosis of errors.

A final advantage is the fact that the R programming language and its packages are open-source. This means that anyone can download it and use the same tools on their data. This also implies that R and its packages can easily be run within virtual environments, either locally or in the cloud, with ease. To this end, I have created a *kinfitr* Docker container, which allows users to download and run a pre-installed version of R, Rstudio and *kinfitr* without needing to install or compile anything on their local machine other than Docker. It is available from the following link: https://hub.docker.com/r/mathesong/kinfitr_docker.

### Usage and validation

The *kinfitr* package has already been used in several scientific papers (Matheson et al. 2017, 2018; Plavén-Sigray et al. 2018; Chen, Goldsmith, and Ogden, n.d.; Stenkrona et al. 2019), and a full validation is currently underway for the consistency of its outcomes compared to other software (Tjerkaski et al., *in prep*). In order to demonstrate basic usage of the *kinfitr* package, as well as to provide a preliminary validation of its outcomes, the package was tested on the sample data included within the package. All of the the relevant models for each dataset were applied for three regions and their estimated binding outcomes are shown in Figure 2. From the figure, it is clear that the estimated parameters using each model are highly correlated with one another. The full reproducible report is included in Supplementary Code.

**Figure 2.**
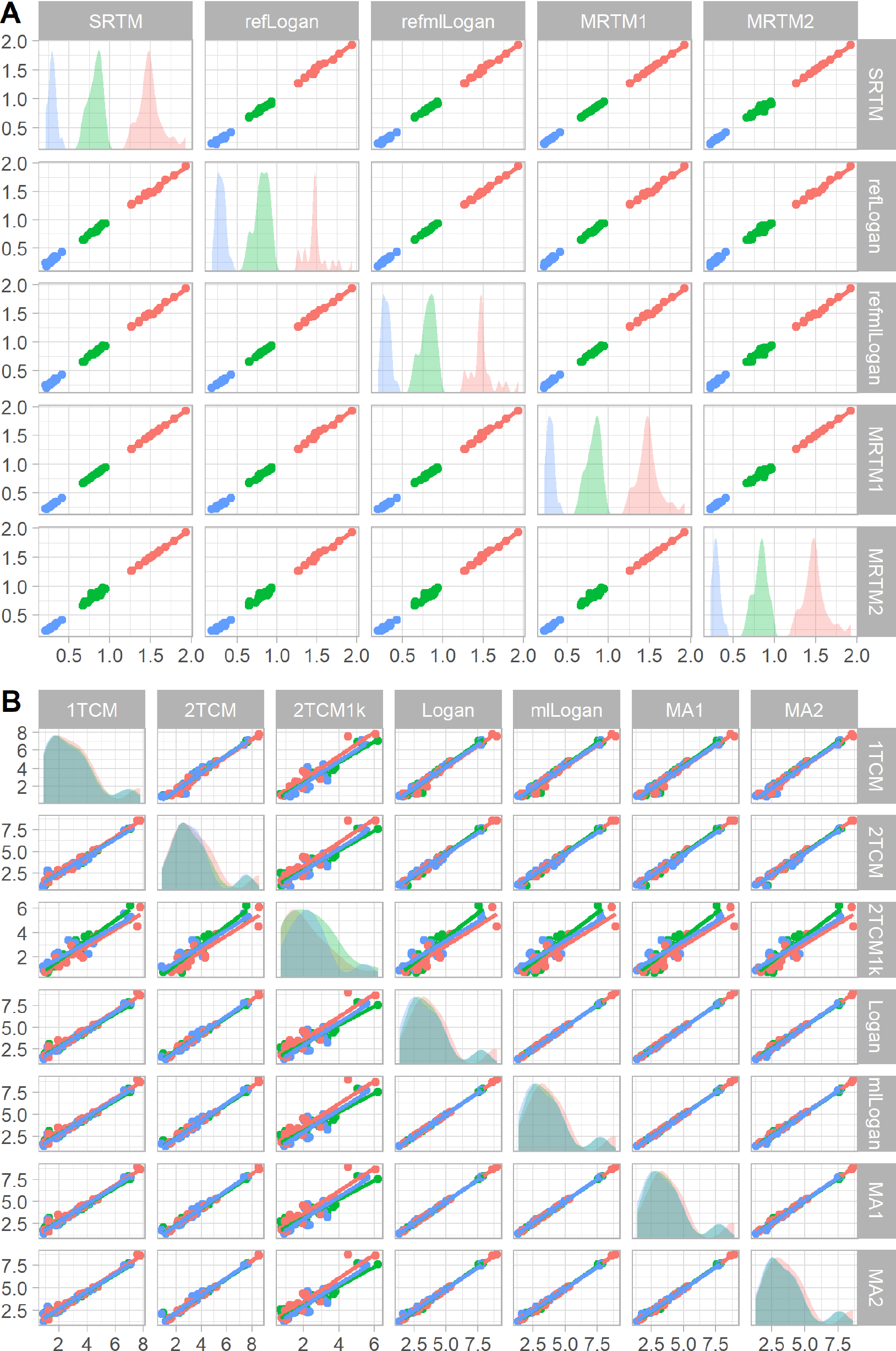
Comparison of primary outcomes between different models which can be applied to the sample data, with regions represented by colour, and regression lines fitted to each combination. A. Comparison of BP ND values obtained using the simref dataset. B. Comparison of V T values obtained using the pbr28 dataset.

## DISCUSSION

I have introduced the *kinfitr* package for analysis of PET TAC data using the R programming language. This package contains tools for processing of PET data following image processing and extraction of time activity curves, until the extraction of binding outcomes from the data and plotting of the model fits. By being situated within the R programming language, this tool can benefit from the extensive collection of other functions and packages within R, as well as the numerous tools for reproducibility, including reproducible reporting and the use of pre-installed virtual computing environments.

The great expense and technical difficulty of PET, especially when blood data is also collected, as well as the fact that participants are injected with harmful radioactivity, makes it imperative that the resulting data is used in an optimal fashion. The *kinfitr* package makes it possible to make better use of PET data, by providing researchers with access to a wide variety of kinetic models, and allows the results of this modelling to be effectively, and transparently communicated in reproducible reports.

This package additionally makes it easier for multi-centre collaborative projects to harmonise their data modelling procedures, as all analysis procedures and instructions are contained within the code which can be shared between centres. By its use of BIDS PET structure for blood data, this means that this complicated data originating from numerous different sources can be quickly and uniformly read and analysed.

In summary, it is hoped that this package will help a researchers to perform PET modelling in a more reproducible fashion, and to prioritise accuracy and transparency to a greater extent in their research. Furthermore, by this project being open-source and hosted on GitHub, other users will also be able to add additional tools and models to the software through pull requests, which can be merged to improve the software package for everyone using it.

## ACKNOWLEDGEMENTS

I gratefully thank the members of the PET group at Karolinska Institutet for their insightful comments and feedback on *kinfitr* over the past years. Thank you especially to Pontus Plavén-Sigray, Jonathan Tjerkaski, Zsolt Cselényi (KI), Yakuan Chen (Columbia University) for their helpful suggestions, feedback, piloting and bug reports. And thank you to Simon Cervenka for allowing me the freedom to work on developing kinfitr during my studies.

