## Supplementary Code for "kinfitr: Reproducible PET Pharmacokinetic Modelling in R"

### kinfitr: Demonstration and Examples

#### Aims

Here I will demonstrate the use of *kinfitr*. I will restrict these examples to estimating macroparameter estimates of binding for reversible tracers, as this is the most common application of PET kinetic modelling, and because we can do this using the included data.

#### Packages

```
library(kinfitr)
```

```
## Registered S3 methods overwritten by 'ggplot2':
##   method         from
##   [.quosures     rlang
##   c.quosures     rlang
##   print.quosures rlang
```

```
library(tidyverse)
```

```
## -- Attaching packages --------------------------------------------------------------------- tidyvers

## v ggplot2 3.1.1     v purrr   0.3.2
## v tibble  2.1.3     v dplyr   0.8.1
## v tidyr   0.8.3     v stringr 1.4.0
## v readr   1.3.1     v forcats 0.4.0

## -- Conflicts ------------------------------------------------------------------------ tidyverse_confl
## x dplyr::filter() masks stats::filter()
## x dplyr::lag()    masks stats::lag()
```

```
library(knitr)
library(ggcorrplot)
```

```
## Warning: package 'ggcorrplot' was built under R version 3.6.1
```

```
library(cowplot)
```

```
## Warning: package 'cowplot' was built under R version 3.6.1

##
## ********************************************************

## Note: As of version 1.0.0, cowplot does not change the

##   default ggplot2 theme anymore. To recover the previous

##   behavior, execute:
##   theme_set(theme_cowplot())

## ********************************************************
```

```
library(mgcv)
```

```
## Loading required package: nlme

##
## Attaching package: 'nlme'
```

```
## The following object is masked from 'package:dplyr':
##
##     collapse
## This is mgcv 1.8-28. For overview type 'help("mgcv-package")'.
```

```
library(ggforce)
```

```
## Warning: package 'ggforce' was built under R version 3.6.1
```

```
theme_set(theme_light())
```

### Reference Region Models

I will start off with a demonstration of how *kinfitr* can be used to fit reference region models. I will use the dataset included in the package, called *simref*. This dataset is simulated to correspond to data which can be successfully modelled with reversible reference-region approaches, with one-tissue dynamics in all regions.

```
data(simref)
```

Let's have a look at the data:

```
head(simref)
```

```
## # A tibble: 6 x 4
##   Subjname PETNo PET    tacs
##   <chr>    <dbl> <chr>  <list>
## 1 hukw         1 hukw_1 <tibble [38 x 8]>
## 2 ybnx         1 ybnx_1 <tibble [38 x 8]>
## 3 olyl         1 olyl_1 <tibble [38 x 8]>
## 4 rocx         1 rocx_1 <tibble [38 x 8]>
## 5 gbiy         1 gbiy_1 <tibble [38 x 8]>
## 6 xsqz         1 xsqz_1 <tibble [38 x 8]>
```

... and within the nested data under tacs is the following:

```
head(simref$tacs[[1]])
```

```
## # A tibble: 6 x 8
##   Times Reference   ROI1    ROI2   ROI3 Weights StartTime Duration
##   <dbl>     <dbl>  <dbl>   <dbl>  <dbl>   <dbl>     <dbl>    <dbl>
## 1 0         0      0       0      0       0         0        0
## 2 0.567     0.137  0.203   0.0518 0.412   0.712     0.483    0.167
## 3 0.733     0.787  1.41    2.31   3.02    0.734     0.65     0.167
## 4 0.9      14.0   17.2    15.3   14.1     0.825     0.817    0.167
## 5 1.07     34.0   41.5    35.5   35.0     0.901     0.983    0.167
## 6 1.23     46.3   57.5    50.7   48.2     0.941     1.15     0.167
```

#### Fitting the data

##### Fitting a single TAC

First let's fit a single TAC to see how everything works

```
tacdata_1 <- simref$tacs[[1]]

t_tac <- tacdata_1$Times
reftac <- tacdata_1$Reference
roitac <- tacdata_1$ROI1
```

```r
weights <- tacdata_1$Weights

fit1 <- srtm(t_tac, reftac, roitac)
```

Now let's look at the fit

```r
plot(fit1)
```

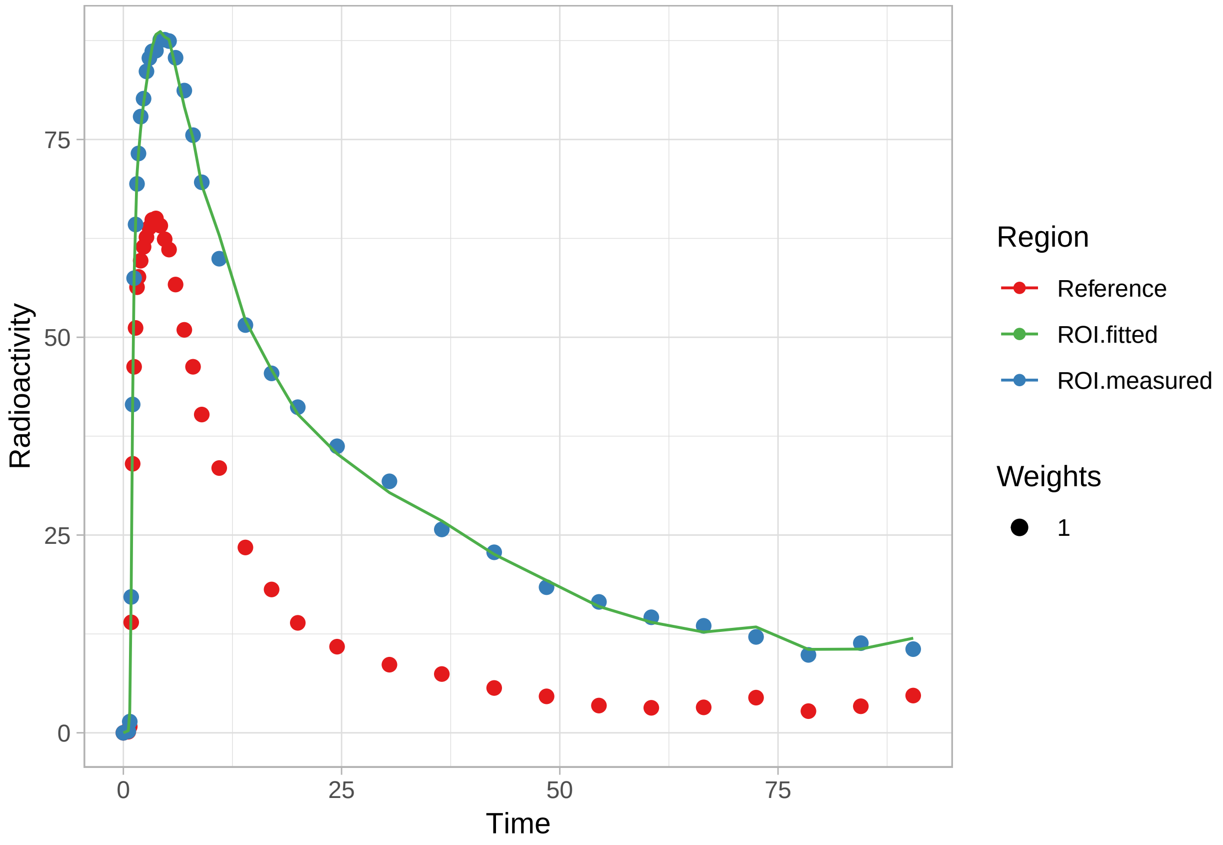


```r
plot_residuals(fit1)
```

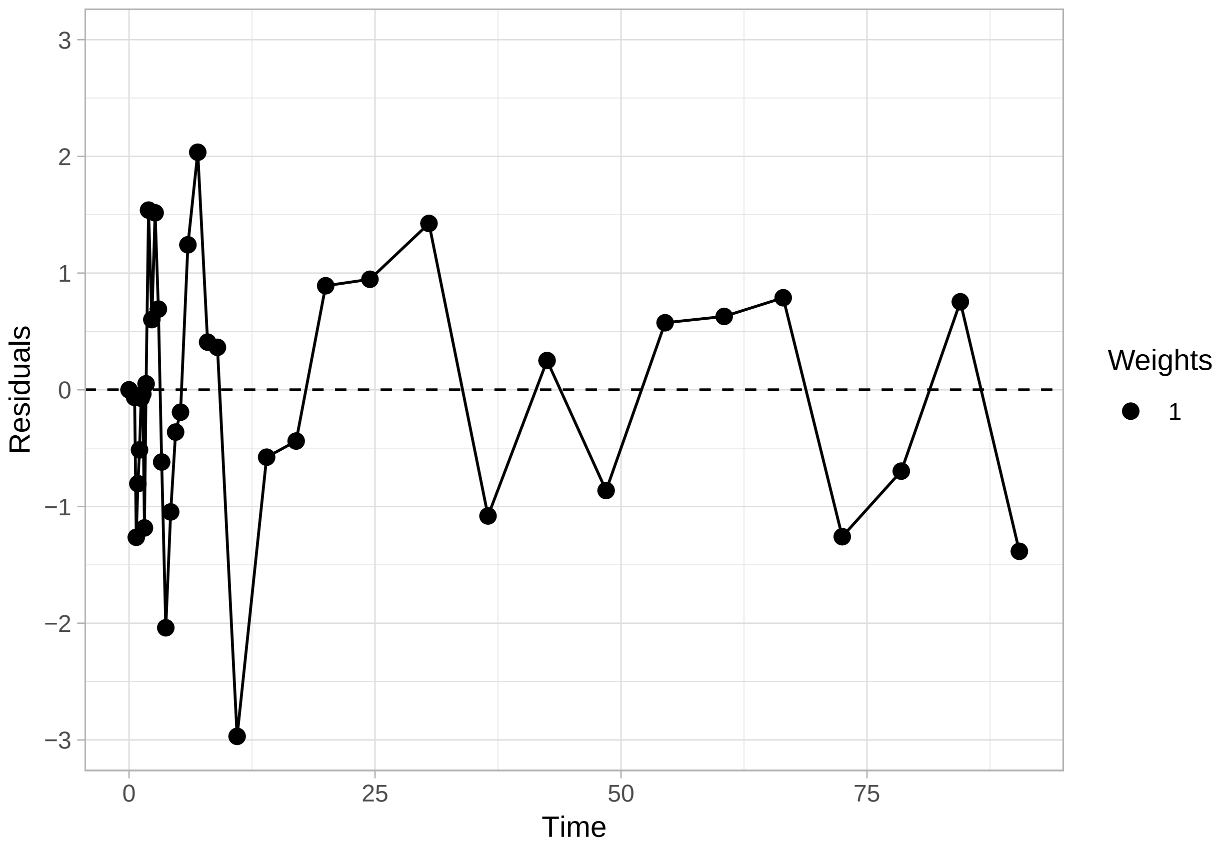


And what is in the fit object?

```
str(fit1)
```

```
## List of 6
##  $ par    :'data.frame': 1 obs. of  3 variables:
##   ..$ R1: num 1.23
##   ..$ k2: num 0.102
##   ..$ bp: num 1.49
##  $ par.se :'data.frame': 1 obs. of  3 variables:
##   ..$ R1.se: num 0.00499
##   ..$ k2.se: num 0.0322
##   ..$ bp.se: num 0.0199
##  $ fit    :List of 6
##   ..$ m       :List of 16
##   .. ..$ resid     :function ()
##   .. ..$ fitted    :function ()
##   .. ..$ formula   :function ()
##   .. ..$ deviance  :function ()
##   .. ..$ lhs       :function ()
##   .. ..$ gradient  :function ()
##   .. ..$ conv      :function ()
##   .. ..$ incr      :function ()
##   .. ..$ setVarying:function (vary = rep(TRUE, length(useParams)))
##   .. ..$ setPars   :function (newPars)
##   .. ..$ getPars   :function ()
##   .. ..$ getAllPars:function ()
```

```
##   .. ..$ getEnv    :function ()
##   .. ..$ trace     :function ()
##   .. ..$ Rmat      :function ()
##   .. ..$ predict   :function (newdata = list(), qr = FALSE)
##   .. ..- attr(*, "class")= chr "nlsModel"
##   ..$ convInfo:List of 5
##   .. ..$ isConv     : logi TRUE
##   .. ..$ finIter    : int 4
##   .. ..$ finTol     : num 1.49e-08
##   .. ..$ stopCode   : int 1
##   .. ..$ stopMessage: chr "Relative error in the sum of squares is at most `ftol'."
##   ..$ data    : symbol modeldata
##   ..$ call    : language minpack.lm::nlsLM(formula = roitac ~ srtm_model(t_tac, reftac, R1,      k2,
##   ..$ weights : num [1:38] 1 1 1 1 1 1 1 1 1 1 ...
##   ..$ control :List of 9
##   .. ..$ ftol   : num 1.49e-08
##   .. ..$ ptol   : num 1.49e-08
##   .. ..$ gtol   : num 0
##   .. ..$ diag   : list()
##   .. ..$ epsfcn : num 0
##   .. ..$ factor : num 100
##   .. ..$ maxfev : int(0)
##   .. ..$ maxiter: num 200
##   .. ..$ nprint : num 0
##   ..- attr(*, "class")= chr "nls"
##  $ weights: num [1:38] 1 1 1 1 1 1 1 1 1 1 ...
##  $ tacs   :'data.frame': 38 obs. of  4 variables:
##   ..$ Time         : num [1:38] 0 0.567 0.733 0.9 1.067 ...
##   ..$ Reference    : num [1:38] 0 0.137 0.787 13.964 34.008 ...
##   ..$ Target       : num [1:38] 0 0.203 1.413 17.179 41.503 ...
##   ..$ Target_fitted: num [1:38] 1.81e-15 2.68e-01 2.68 1.80e+01 4.20e+01 ...
##  $ model  : chr "srtm"
##  - attr(*, "class")= chr [1:2] "srtm" "kinfit"
```

There's a lot.

Of primary importance:

- **par** contains the parameters
- **par.se** contains the standard errors as a proportion of the values
- **fit** contains the actual model fit object, which we can do all sorts of crazy things with
- **weights**, **tacs**, **model** are pretty self-explanatory

The fit object being there means we can do all of the things that R, being a language designed by statisticians, allows us to do easily

```
g <- fit1$fit

coef(g)
```

```
##        R1        k2        bp
## 1.2335455 0.1016237 1.4883389
```

```
resid(g)
```

```
##  [1] -1.814790e-15 -6.503612e-02 -1.263658e+00 -8.041934e-01 -5.149457e-01
##  [6] -7.244154e-02 -3.525757e-02 -1.183420e+00  5.159218e-02  1.539581e+00
```

```
## [11]  6.016639e-01  1.516351e+00  6.913366e-01 -6.188564e-01 -2.039015e+00
## [16] -1.046386e+00 -3.622996e-01 -1.916459e-01  1.242381e+00  2.035547e+00
## [21]  4.086986e-01  3.637660e-01 -2.969245e+00 -5.773616e-01 -4.389941e-01
## [26]  8.916004e-01  9.471013e-01  1.425524e+00 -1.081132e+00  2.508164e-01
## [31] -8.628823e-01  5.746597e-01  6.287208e-01  7.888887e-01 -1.258560e+00
## [36] -6.968996e-01  7.543217e-01 -1.383989e+00
## attr(,"label")
## [1] "Residuals"
```

```
vcov(g)
```

```
##               R1            k2            bp
## R1  3.783235e-05 -1.085198e-05  3.109032e-05
## k2 -1.085198e-05  1.073623e-05 -5.748971e-05
## bp  3.109032e-05 -5.748971e-05  8.795270e-04
```

```
AIC(g)
```

```
## [1] 119.7931
```

```
BIC(g)
```

```
## [1] 126.3434
```

#### Multiple TACs

Now let's fit a model to more TACs, one from each individual. I'll also use a different model: in this case MRTM1.

```
simref <- simref %>%
  group_by(Subjname, PETNo) %>%
  mutate(MRTM1fit = map(tacs, ~mrtm1(t_tac = .x$Times, reftac = .x$Reference,
                                     roitac = .x$ROI1,
                                     weights = .x$Weights))) %>%
  ungroup()
```

Now let's look at the distribution of our $BP_{ND}$ values

```
simref <- simref %>%
  mutate(bp_MRTM1 = map_dbl(MRTM1fit, c("par", "bp")))

ggplot(simref, aes(x=bp_MRTM1)) +
  geom_histogram(fill="grey", colour="black")
```

```
## `stat_bin()` using `bins = 30`. Pick better value with `binwidth`.
```

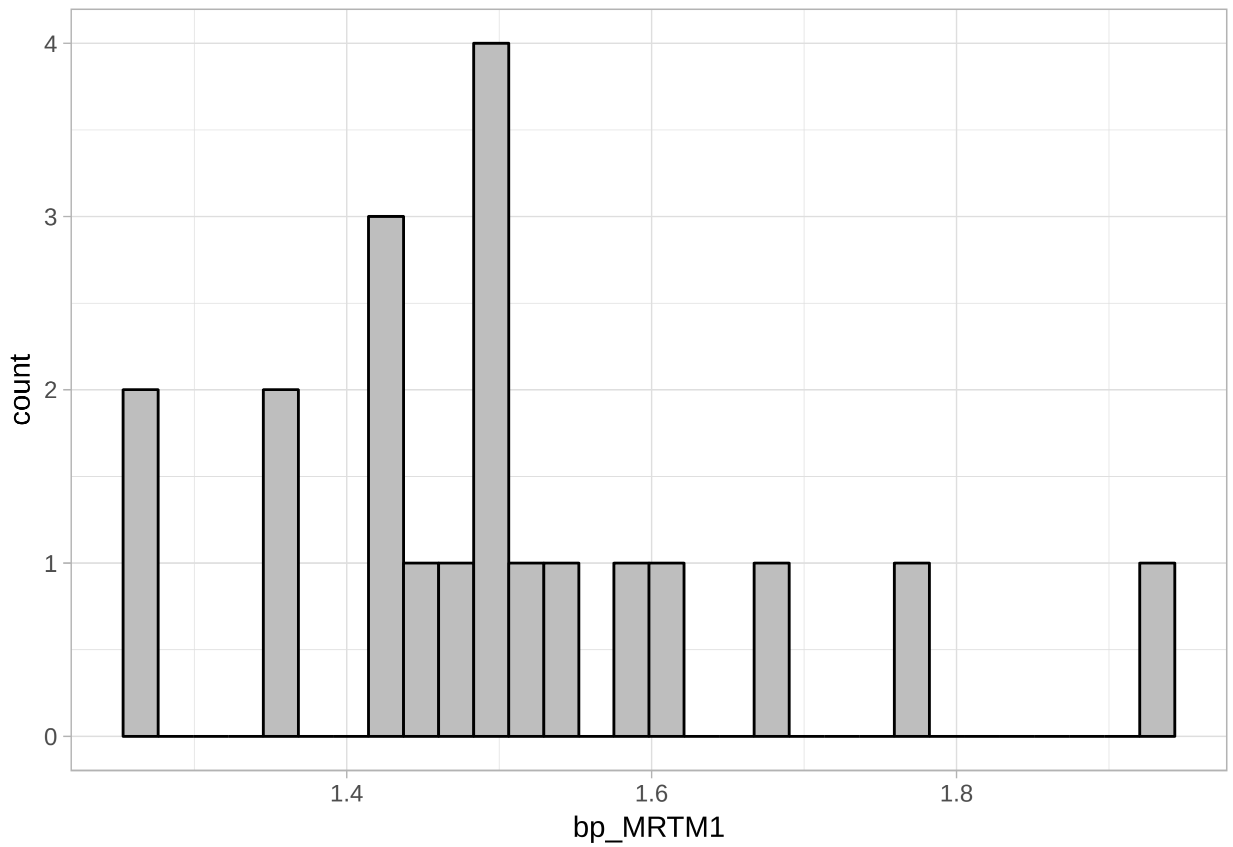


#### Multiple Regions from Multiple Individuals

Now, let's fit all the regions, and use multiple models. First, though, we need to account for some extra requirements of some other models.

##### k2prime

One requirement of several models is that a k2prime value be specified. This can be done by fitting this value using one of the models which fits it. k2prime can be assessed using SRTM indirectly, using R1 and k2, or directly using MRTM1. It should also be stable across the brain, since it is a function of the reference region, and therefore it can be fitted for one region which we have already done above. Let's first extract the k2prime values into their own column first.

```
simref <- simref %>%
  mutate(k2prime = map_dbl(MRTM1fit, c("par", "k2prime")))

ggplot(simref, aes(x=k2prime)) +
  geom_histogram(fill="grey", colour="black")
```

```
## `stat_bin()` using `bins = 30`. Pick better value with `binwidth`.
```

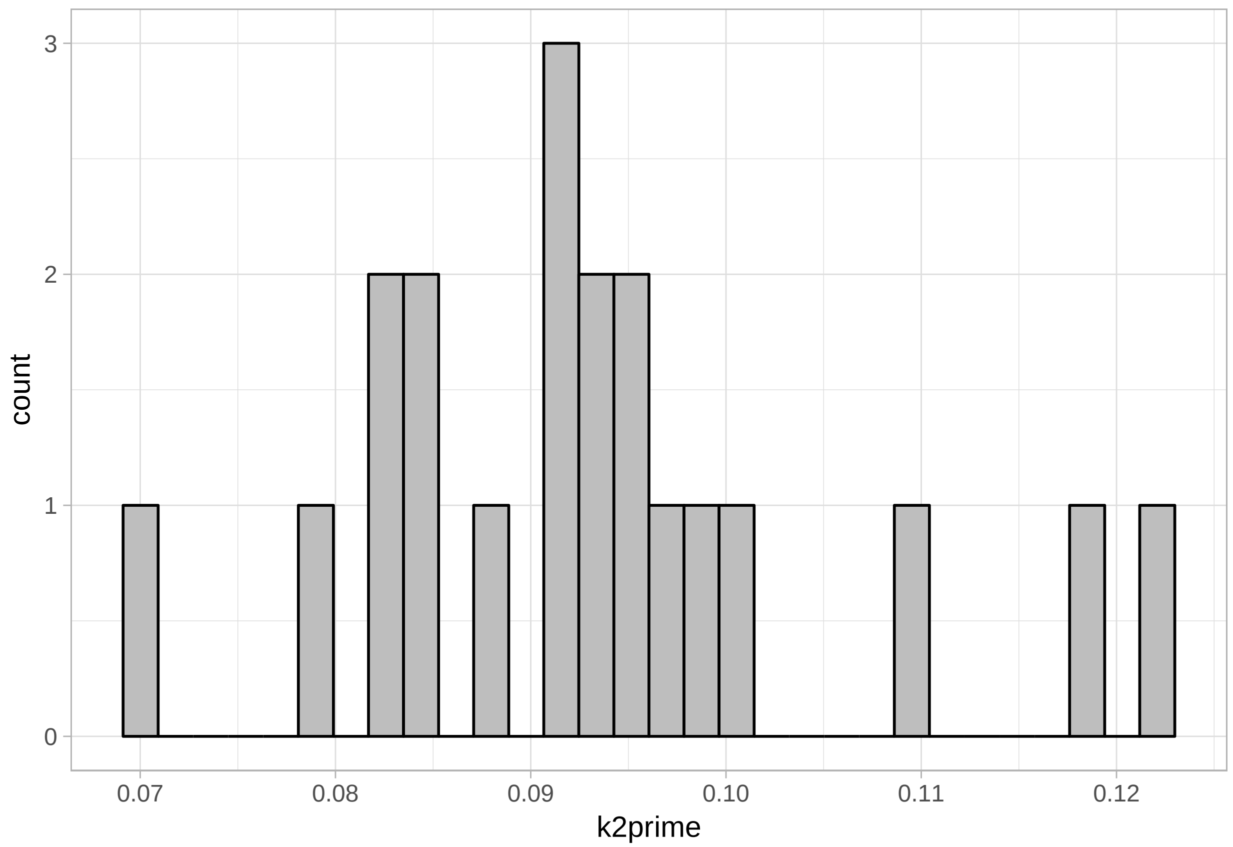


**t star**

Just about all linearised models used in PET have a t* value. This is the point at which the linear fit becomes linear, and the value provided counts the number of frames from the end of the end of the PET meausrement to be included.

In MRTM1 and MRTM2, the t* value is necessary only if the dynamics do not correspond to one-tissue compartment dynamics, which the simulated values specifically do. We will, however, need to identify a t* value for the non-invasive Logan plot and the multilinear non-ínvasive Logan plot.

Let's take a look at a few first for the non-invasive Logan plot

```
set.seed(12345)

subs <- sample(1:nrow(simref), size = 3, replace = F)

s = subs[1]

td <- simref$tacs[[s]]
refLogan_tstar(t_tac = td$Times, reftac = td$Reference, lowroi = td$ROI3,
               medroi = td$ROI2, highroi = td$ROI1,
               k2prime = simref$k2prime[s])
```

```
## Warning: Removed 1 rows containing missing values (geom_point).

## Warning: Removed 1 rows containing missing values (geom_point).

## Warning: Removed 1 rows containing missing values (geom_point).
```

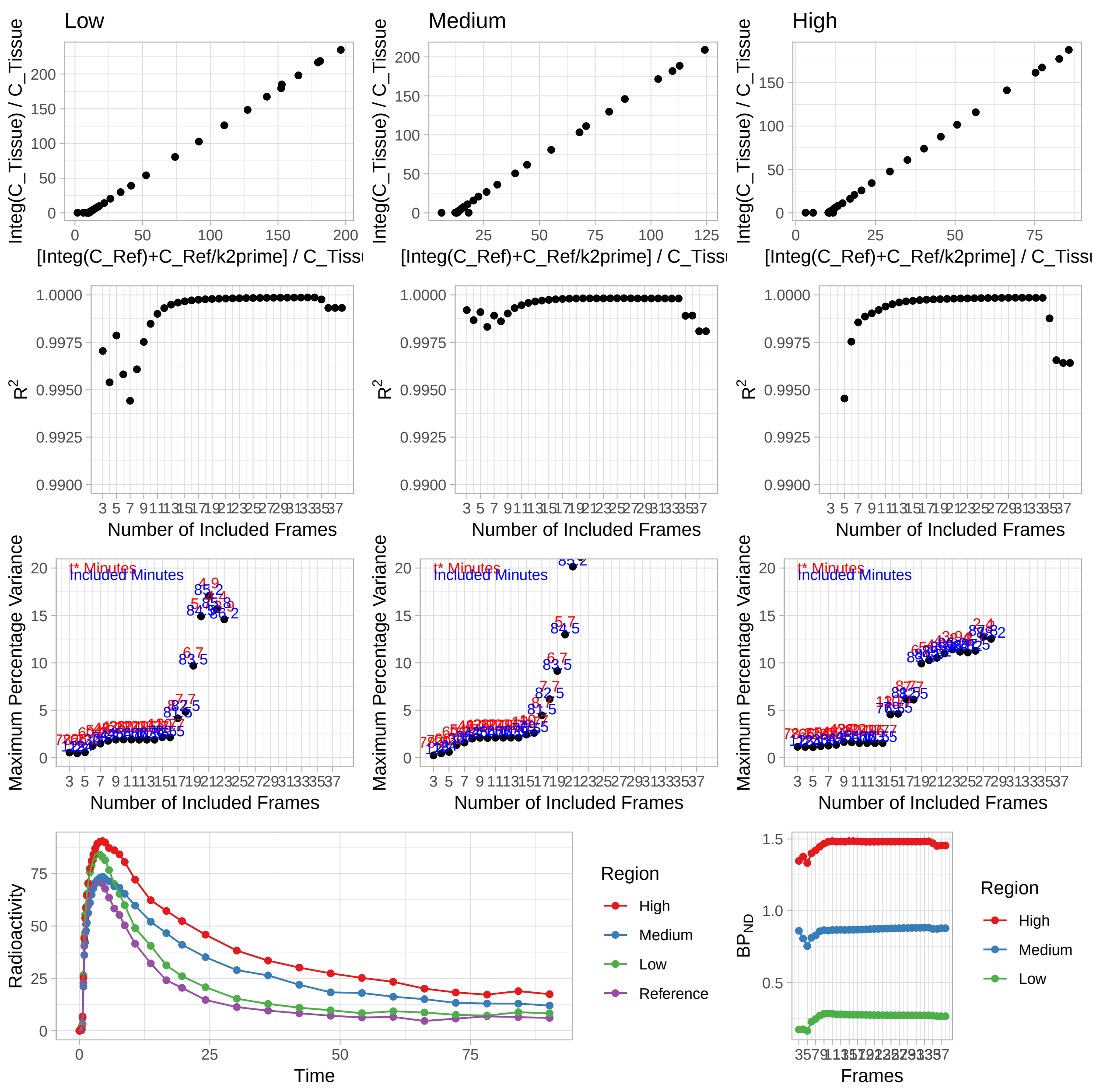


```r
s = subs[2]

td <- simref$tacs[[s]]
refLogan_tstar(t_tac = td$Times, reftac = td$Reference, lowroi = td$ROI3,
               medroi = td$ROI2, highroi = td$ROI1,
               k2prime = simref$k2prime[s])
```

```
## Warning: Removed 1 rows containing missing values (geom_point).

## Warning: Removed 1 rows containing missing values (geom_point).

## Warning: Removed 1 rows containing missing values (geom_point).
```

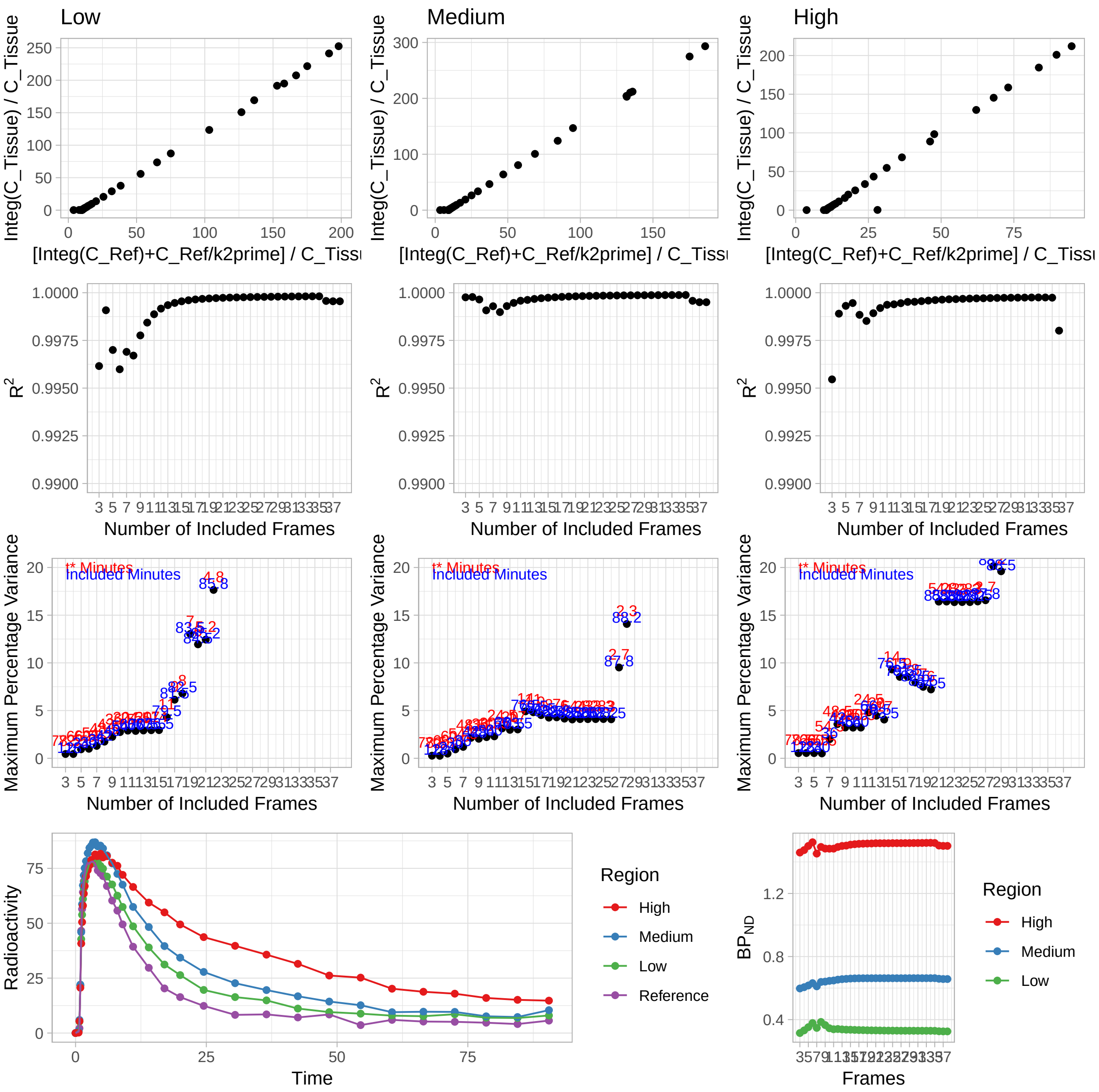


```r
s = subs[3]

td <- simref$tacs[[s]]
refLogan_tstar(t_tac = td$Times, reftac = td$Reference, lowroi = td$ROI3,
               medroi = td$ROI2, highroi = td$ROI1,
               k2prime = simref$k2prime[s])
```

```
## Warning: Removed 1 rows containing missing values (geom_point).

## Warning: Removed 1 rows containing missing values (geom_point).

## Warning: Removed 1 rows containing missing values (geom_point).
```

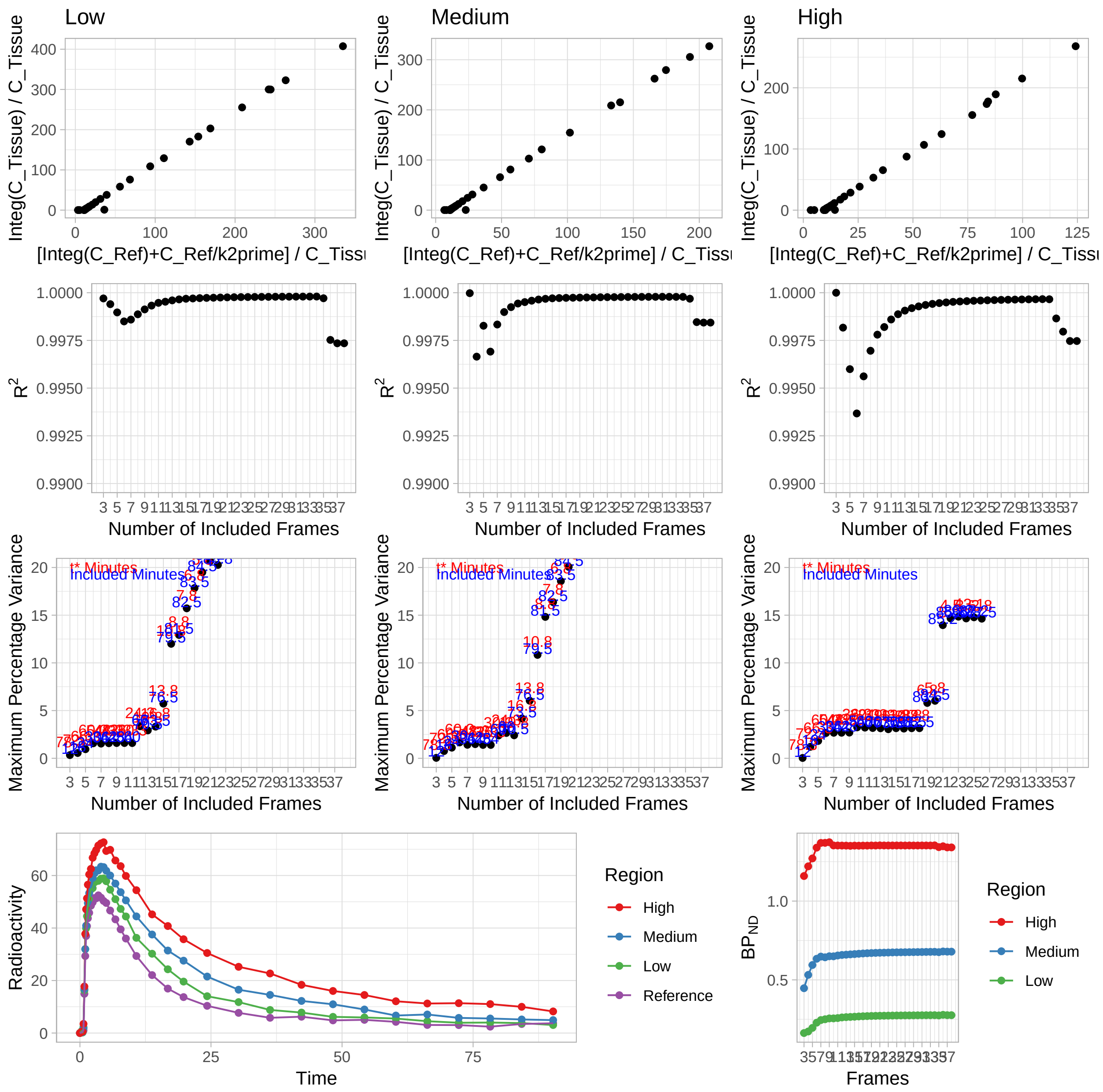


We can see the following in general:

- The $R^2$ values are pretty high after around 10 frames
- The maximum percentage variance increases quickly after around the same point, though it is earlier and later for different individuals and different regions. But 10 seems to be fine for all of them.
- Also, after about 10 frames, the $BP_{ND}$ values are quite stable (bottom right)

So, we will use 10 frames for the t*. Let's check if we come to other conclusions for the multilinear non-invasive Logan plot

```
set.seed(42)

subs <- sample(1:nrow(simref), size = 3, replace = F)

s = subs[1]
```

```r
td <- simref$tacs[[s]]
refmlLogan_tstar(t_tac = td$Times, reftac = td$Reference, lowroi = td$ROI3,
                 medroi = td$ROI2, highroi = td$ROI1,
                 k2prime = simref$k2prime[s])
```

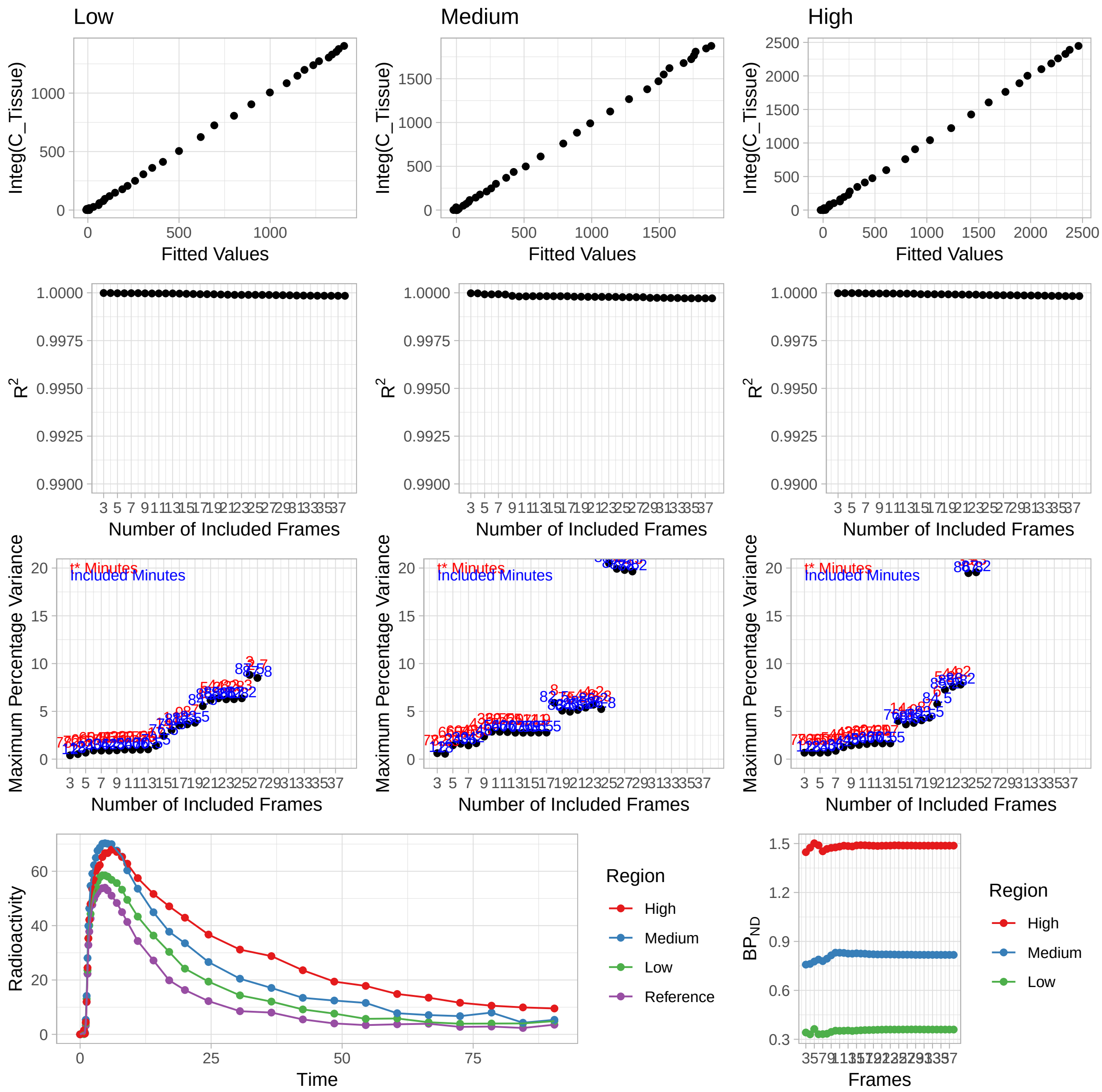


```r
s = subs[2]

td <- simref$tacs[[s]]
refmlLogan_tstar(t_tac = td$Times, reftac = td$Reference, lowroi = td$ROI3,
                 medroi = td$ROI2, highroi = td$ROI1,
                 k2prime = simref$k2prime[s])
```

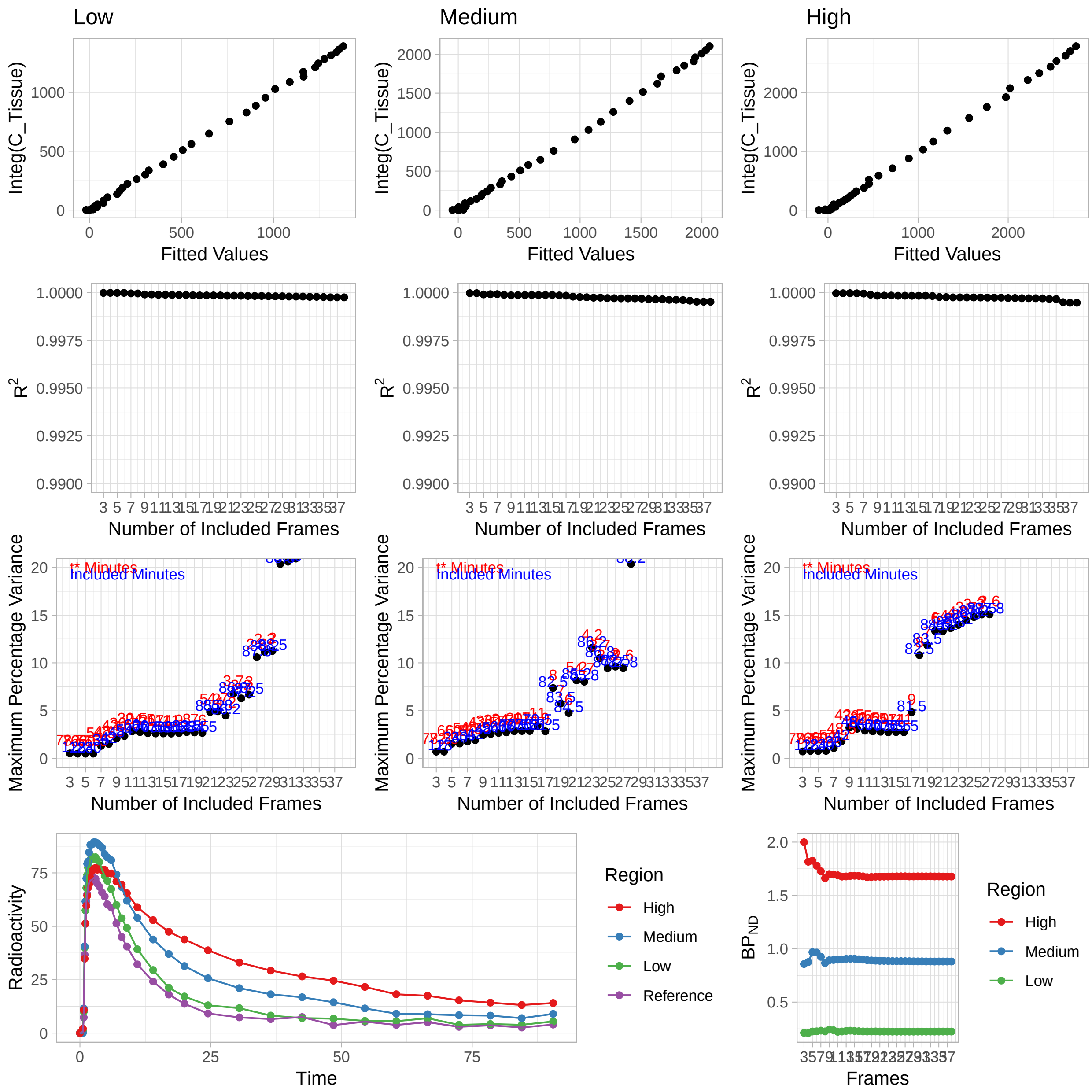


```
s = subs[3]

td <- simref$tacs[[s]]
refmlLogan_tstar(t_tac = td$Times, reftac = td$Reference, lowroi = td$ROI3,
                 medroi = td$ROI2, highroi = td$ROI1,
                 k2prime = simref$k2prime[s])
```

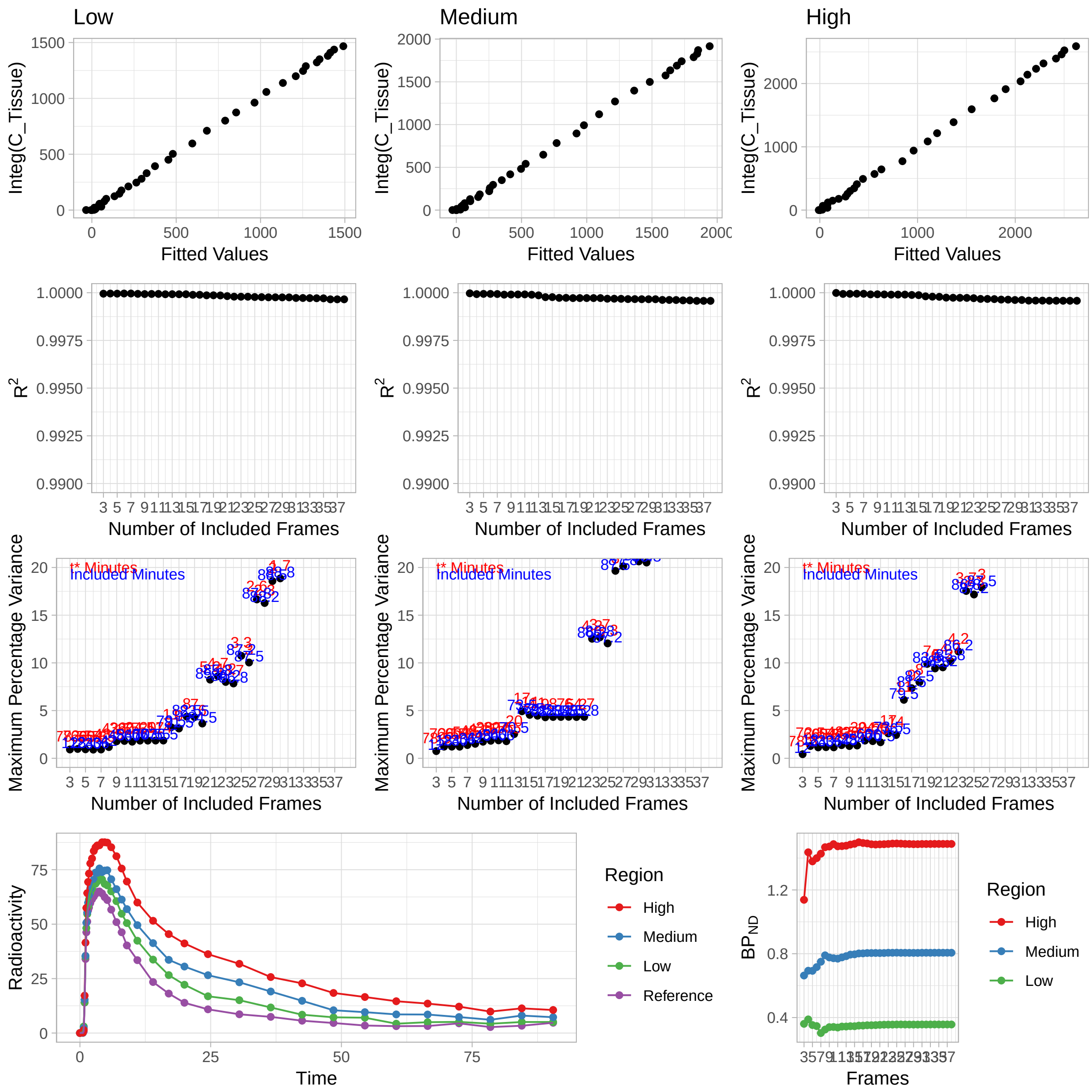


This looks a bit more stable, Let's go for 13 in this case.

##### Model fitting

Now, we have extracted estimated k2prime values, and selected t* values. We are ready to fit most models.

First, we rearrange the data a little bit, so that it is chunked at the level of ROIs within regions, as opposed to individuals. Currently the data looks as follows:

```
simref <- simref %>%
  select(-MRTM1fit, -bp_MRTM1)

simref
```

```
## # A tibble: 20 x 5
##    Subjname PETNo PET    tacs                 k2prime
```

```
##     <chr>     <dbl> <chr>  <list>              <dbl>
##   1 hukw          1 hukw_1 <tibble [38 x 8]>  0.0825
##   2 ybnx          1 ybnx_1 <tibble [38 x 8]>  0.119
##   3 olyl          1 olyl_1 <tibble [38 x 8]>  0.0972
##   4 rocx          1 rocx_1 <tibble [38 x 8]>  0.0934
##   5 gbiy          1 gbiy_1 <tibble [38 x 8]>  0.101
##   6 xsqz          1 xsqz_1 <tibble [38 x 8]>  0.0923
##   7 rsop          1 rsop_1 <tibble [38 x 8]>  0.0910
##   8 hdzx          1 hdzx_1 <tibble [38 x 8]>  0.0990
##   9 ruam          1 ruam_1 <tibble [38 x 8]>  0.0782
##  10 tfig          1 tfig_1 <tibble [38 x 8]>  0.0949
##  11 dkkj          1 dkkj_1 <tibble [38 x 8]>  0.0911
##  12 ddgm          1 ddgm_1 <tibble [38 x 8]>  0.0851
##  13 gwbl          1 gwbl_1 <tibble [38 x 8]>  0.122
##  14 udof          1 udof_1 <tibble [38 x 8]>  0.0872
##  15 dtxj          1 dtxj_1 <tibble [38 x 8]>  0.0832
##  16 rcjh          1 rcjh_1 <tibble [38 x 8]>  0.0844
##  17 vlvv          1 vlvv_1 <tibble [38 x 8]>  0.0943
##  18 ultq          1 ultq_1 <tibble [38 x 8]>  0.0702
##  19 samf          1 samf_1 <tibble [38 x 8]>  0.110
##  20 jpjc          1 jpjc_1 <tibble [38 x 8]>  0.0928
```

Next, we want to chunk at a different level. Instead of individuals, we want to chunk at regions of each individual.

```
simref_long <- simref %>%
  unnest() %>%
  select(-StartTime, -Duration, -PET)

simref_long
```

```
## # A tibble: 760 x 9
##     Subjname PETNo k2prime Times Reference   ROI1    ROI2   ROI3 Weights
##     <chr>    <dbl>   <dbl> <dbl>     <dbl>  <dbl>   <dbl>  <dbl>   <dbl>
##   1 hukw         1  0.0825 0         0      0       0      0       0
##   2 hukw         1  0.0825 0.567     0.137  0.203   0.0518 0.412   0.712
##   3 hukw         1  0.0825 0.733     0.787  1.41    2.31   3.02    0.734
##   4 hukw         1  0.0825 0.9      14.0   17.2    15.3   14.1     0.825
##   5 hukw         1  0.0825 1.07     34.0   41.5    35.5   35.0     0.901
##   6 hukw         1  0.0825 1.23     46.3   57.5    50.7   48.2     0.941
##   7 hukw         1  0.0825 1.4      51.2   64.3    55.4   54.6     0.958
##   8 hukw         1  0.0825 1.57     56.3   69.4    59.9   58.2     0.970
##   9 hukw         1  0.0825 1.73     57.6   73.2    63.5   61.4     0.979
##  10 hukw         1  0.0825 1.98     59.7   77.9    66.4   64.5     0.988
## # ... with 750 more rows
```

And now we make it into a very long format (i.e. 760 rows to 2280 rows).

```
simref_long <- simref_long %>%
  gather(key = Region, value = TAC, -Times, -Weights,
         -Subjname, -PETNo, -k2prime, -Reference)

simref_long
```

```
## # A tibble: 2,280 x 8
##     Subjname PETNo k2prime Times Reference Weights Region    TAC
```

```
##    <chr>    <dbl>   <dbl> <dbl>     <dbl>   <dbl> <chr>    <dbl>
##  1 hukw         1  0.0825 0         0       0     ROI1     0
##  2 hukw         1  0.0825 0.567     0.137   0.712 ROI1     0.203
##  3 hukw         1  0.0825 0.733     0.787   0.734 ROI1     1.41
##  4 hukw         1  0.0825 0.9      14.0     0.825 ROI1    17.2
##  5 hukw         1  0.0825 1.07     34.0     0.901 ROI1    41.5
##  6 hukw         1  0.0825 1.23     46.3     0.941 ROI1    57.5
##  7 hukw         1  0.0825 1.4      51.2     0.958 ROI1    64.3
##  8 hukw         1  0.0825 1.57     56.3     0.970 ROI1    69.4
##  9 hukw         1  0.0825 1.73     57.6     0.979 ROI1    73.2
## 10 hukw         1  0.0825 1.98     59.7     0.988 ROI1    77.9
## # ... with 2,270 more rows
```

And now we nest it again, with our desired level of chunking.

```
simref_long <- simref_long %>%
  group_by(Subjname, PETNo, k2prime, Region) %>%
  nest(.key = "tacdata")

simref_long
```

```
## # A tibble: 60 x 5
##    Subjname PETNo k2prime Region tacdata
##    <chr>    <dbl>   <dbl> <chr>  <list>
##  1 hukw         1  0.0825 ROI1   <tibble [38 x 4]>
##  2 ybnx         1  0.119  ROI1   <tibble [38 x 4]>
##  3 olyl         1  0.0972 ROI1   <tibble [38 x 4]>
##  4 rocx         1  0.0934 ROI1   <tibble [38 x 4]>
##  5 gbiy         1  0.101  ROI1   <tibble [38 x 4]>
##  6 xsqz         1  0.0923 ROI1   <tibble [38 x 4]>
##  7 rsop         1  0.0910 ROI1   <tibble [38 x 4]>
##  8 hdzx         1  0.0990 ROI1   <tibble [38 x 4]>
##  9 ruam         1  0.0782 ROI1   <tibble [38 x 4]>
## 10 tfig         1  0.0949 ROI1   <tibble [38 x 4]>
## # ... with 50 more rows
```

And within each chunk is the following:

```
simref_long$tacdata[[1]]
```

```
## # A tibble: 38 x 4
##    Times Reference Weights    TAC
##    <dbl>     <dbl>   <dbl>  <dbl>
##  1 0         0       0      0
##  2 0.567     0.137   0.712  0.203
##  3 0.733     0.787   0.734  1.41
##  4 0.9      14.0     0.825 17.2
##  5 1.07     34.0     0.901 41.5
##  6 1.23     46.3     0.941 57.5
##  7 1.4      51.2     0.958 64.3
##  8 1.57     56.3     0.970 69.4
##  9 1.73     57.6     0.979 73.2
## 10 1.98     59.7     0.988 77.9
## # ... with 28 more rows
```

Now we can proceed to fitting models to all the regions.

```r
simref_long <- simref_long %>%
  group_by(Subjname, PETNo, Region) %>%

  # SRTM
  mutate(SRTM = map(tacdata,
                        ~srtm(t_tac=.x$Times, reftac = .x$Reference,
                              roitac = .x$TAC,
                              weights = .x$Weights)),
         SRTM = map_dbl(SRTM, c("par", "bp"))) %>%

  # MRTM1
  mutate(MRTM1 = map(tacdata,
                        ~mrtm1(t_tac=.x$Times, reftac = .x$Reference,
                              roitac = .x$TAC,
                              weights = .x$Weights)),
         MRTM1 = map_dbl(MRTM1, c("par", "bp"))) %>%

  # MRTM2
  mutate(MRTM2 = map2(tacdata, k2prime,
                        ~mrtm2(t_tac=.x$Times, reftac = .x$Reference,
                              roitac = .x$TAC, k2prime = .y,
                              weights = .x$Weights)),
         MRTM2 = map_dbl(MRTM2, c("par", "bp"))) %>%

  # refLogan
  mutate(refLogan = map2(tacdata, k2prime,
                        ~refLogan(t_tac=.x$Times, reftac = .x$Reference,
                              roitac = .x$TAC, k2prime = .y,
                              weights = .x$Weights, tstarIncludedFrames = 10)),
         refLogan = map_dbl(refLogan, c("par", "bp"))) %>%

  # refmlLogan
  mutate(refmlLogan = map2(tacdata, k2prime,
                        ~refmlLogan(t_tac=.x$Times, reftac = .x$Reference,
                              roitac = .x$TAC, k2prime = .y,
                              weights = .x$Weights, tstarIncludedFrames = 10)),
         refmlLogan = map_dbl(refmlLogan, c("par", "bp"))) %>%
  ungroup()
```

#### Comparisons between Models

Now let's check the correlations between the estimated $BP_{ND}$ values.

```r
simref_bpvals <- simref_long %>%
  ungroup() %>%
  select(SRTM, refLogan, refmlLogan, MRTM1, MRTM2)

ref_corrplot <- ggcorrplot(cor(simref_bpvals), lab=T,
           colors = c("#6D9EC1", "white", "#E46726"),
           digits = 3, show.diag = F)

print(ref_corrplot)
```

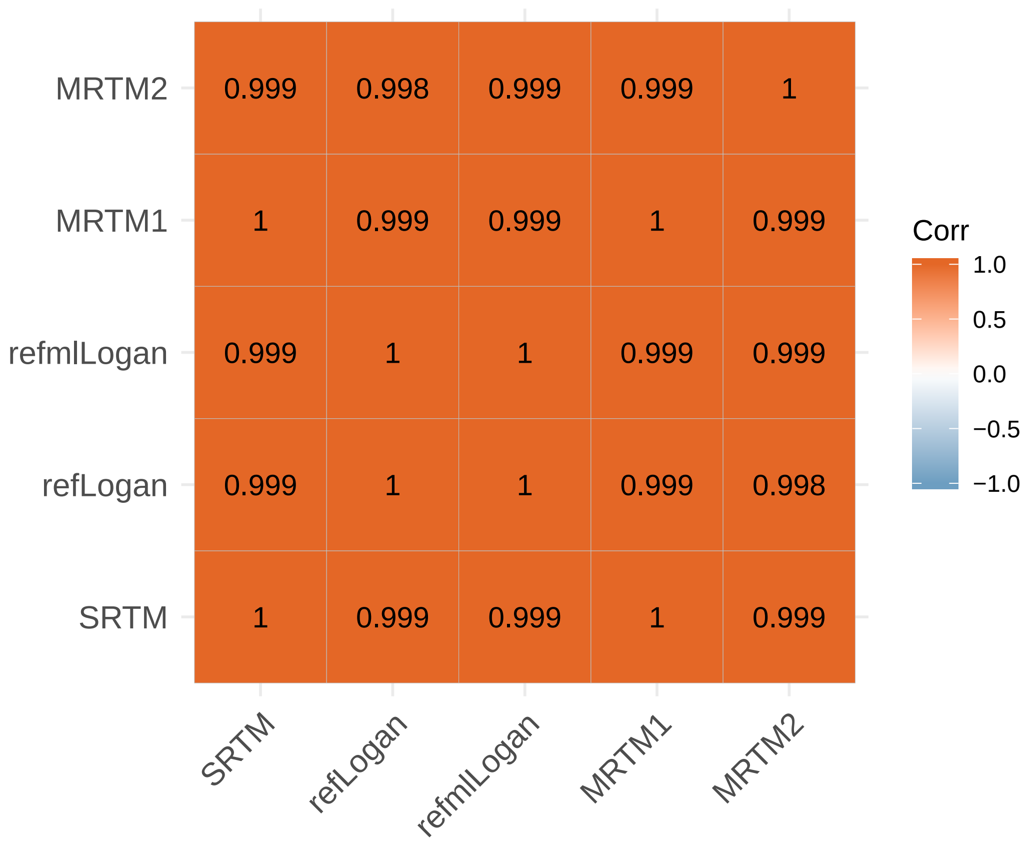


#### Arterial Input Models

Now we will work with some data for which there is no reference region. This is data collected using [11C]PBR28, which binds to the 18kDa translocator protein, which is found throughout the brain, meaning that there is no reference region.

##### Blood Processing

The first, complicated aspect of working with data for which we must use the arterial plasma as a reference is that this data is collected through a parallel data collection, and is stored separately. Most of this issue is resolved by the introduction of BIDS for PET, which provides a JSON sidecar file along with every image file, and the sidecar contains all the information which is required in addition to the image file. kinfitr is able to automatically read these sidecar files, and create a *blooddata* object with them. Within the *pbr28* data within *kinfitr* I have put together the information which would be available within the PET BIDS JSON sidecar file for each measurement in a column called *jsondata*, which can be loaded directly as if it were a JSON file. Here, I will create blooddata objects out of them all.

```
data(pbr28)

pbr28 <- pbr28 %>%
  select(-procblood, -input, -Genotype, -injRad)

head(pbr28)
```

```
## # A tibble: 6 x 5
##   PET    Subjname PETNo tacs                jsondata
```

```
##   <chr>  <chr>    <dbl> <list>             <list>
## 1 cgyu_1 cgyu         1 <tibble [38 x 10]> <list [6]>
## 2 cgyu_2 cgyu         2 <tibble [38 x 10]> <list [6]>
## 3 flfp_1 flfp         1 <tibble [38 x 10]> <list [6]>
## 4 flfp_2 flfp         2 <tibble [38 x 10]> <list [6]>
## 5 jdcs_1 jdcs         1 <tibble [38 x 10]> <list [6]>
## 6 jdcs_2 jdcs         2 <tibble [38 x 10]> <list [6]>
```

Now we can create a *blooddata* object from each.

```
pbr28 <- pbr28 %>%
  mutate(blooddata = map(jsondata, create_blooddata_bids))
```

##### Dispersion Correction

The first step is dispersion correction of blood samples collected through the automatic system, for dispersion as the blood travels through the plastic tubing. This is corrected for using the dispersion constant (included in the BIDS sidecar). All functions which can be used on the *blooddata* objects are prefixed with *bd_*, in this case *bd_blood_dispcor*.

```
pbr28 <- pbr28 %>%
  mutate(blooddata = map(blooddata, ~bd_blood_dispcor(.x, smooth_iterations = 2)))
```

In this case, we first perform dispersion correction, and then smooth the data slightly.

##### Modelling the Blood

Let's have a look at a few blood curves

```
plot(pbr28$blooddata[[1]])
```

```
## Warning: Removed 1 rows containing missing values (geom_path).
```

```
## Warning: Removed 1 rows containing missing values (geom_path).
```

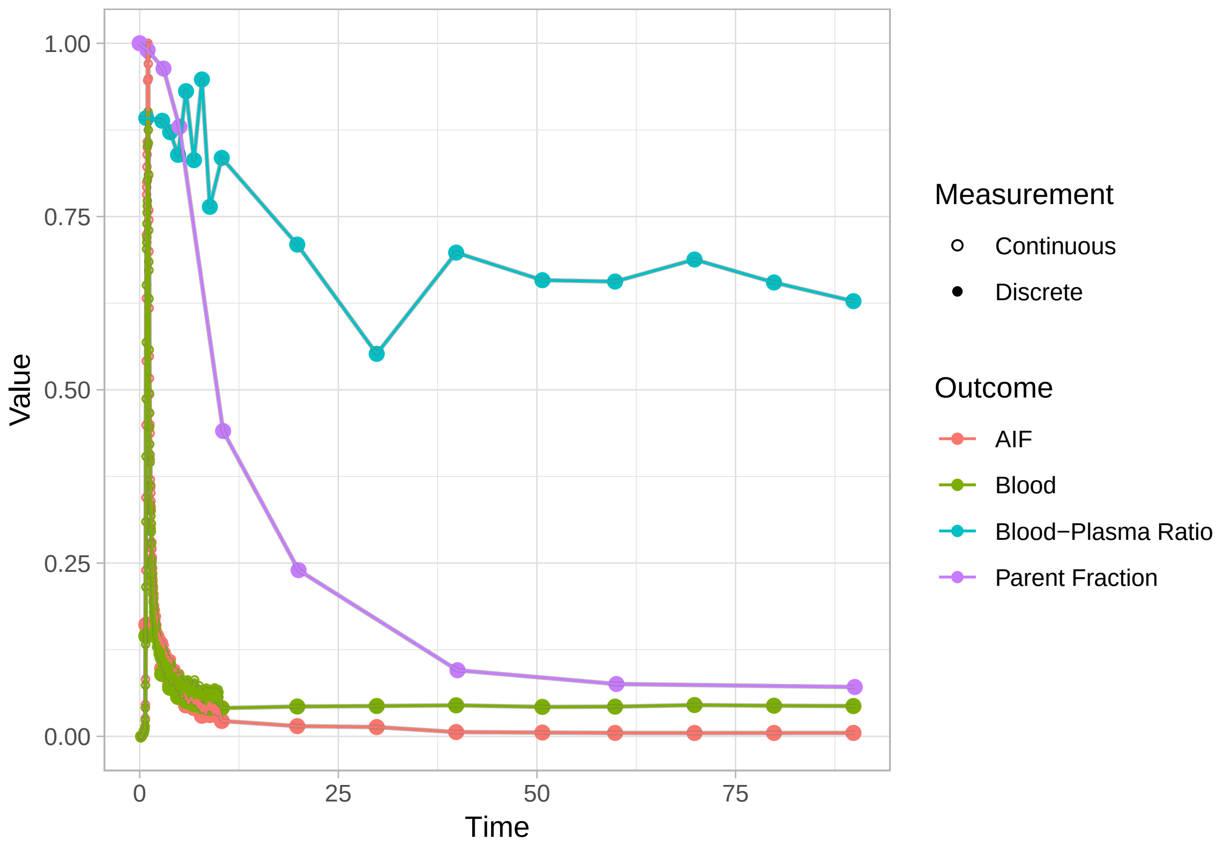


```
plot(pbr28$blooddata[[2]])
```

```
## Warning: Removed 1 rows containing missing values (geom_path).

## Warning: Removed 1 rows containing missing values (geom_path).
```

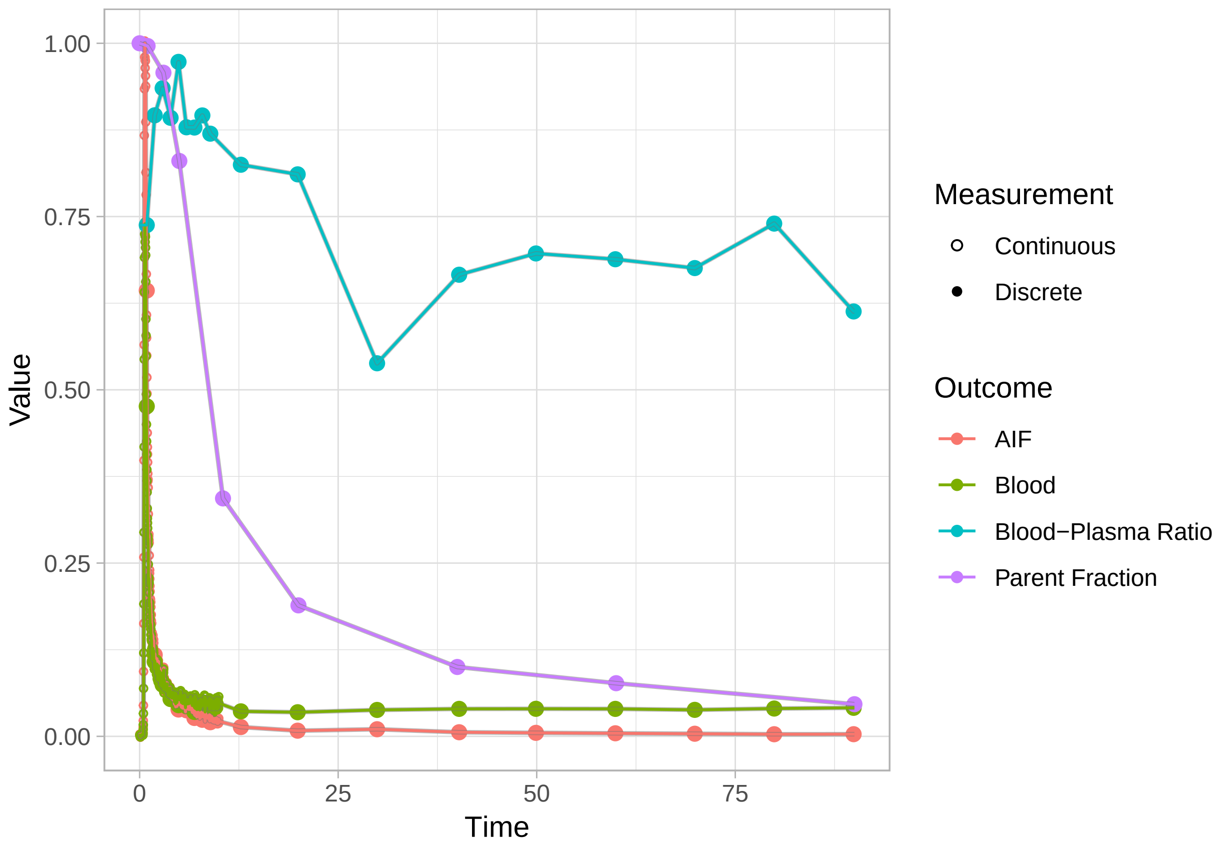


```
plot(pbr28$blooddata[[3]])
```

```
## Warning: Removed 1 rows containing missing values (geom_path).
```

```
## Warning: Removed 1 rows containing missing values (geom_path).
```

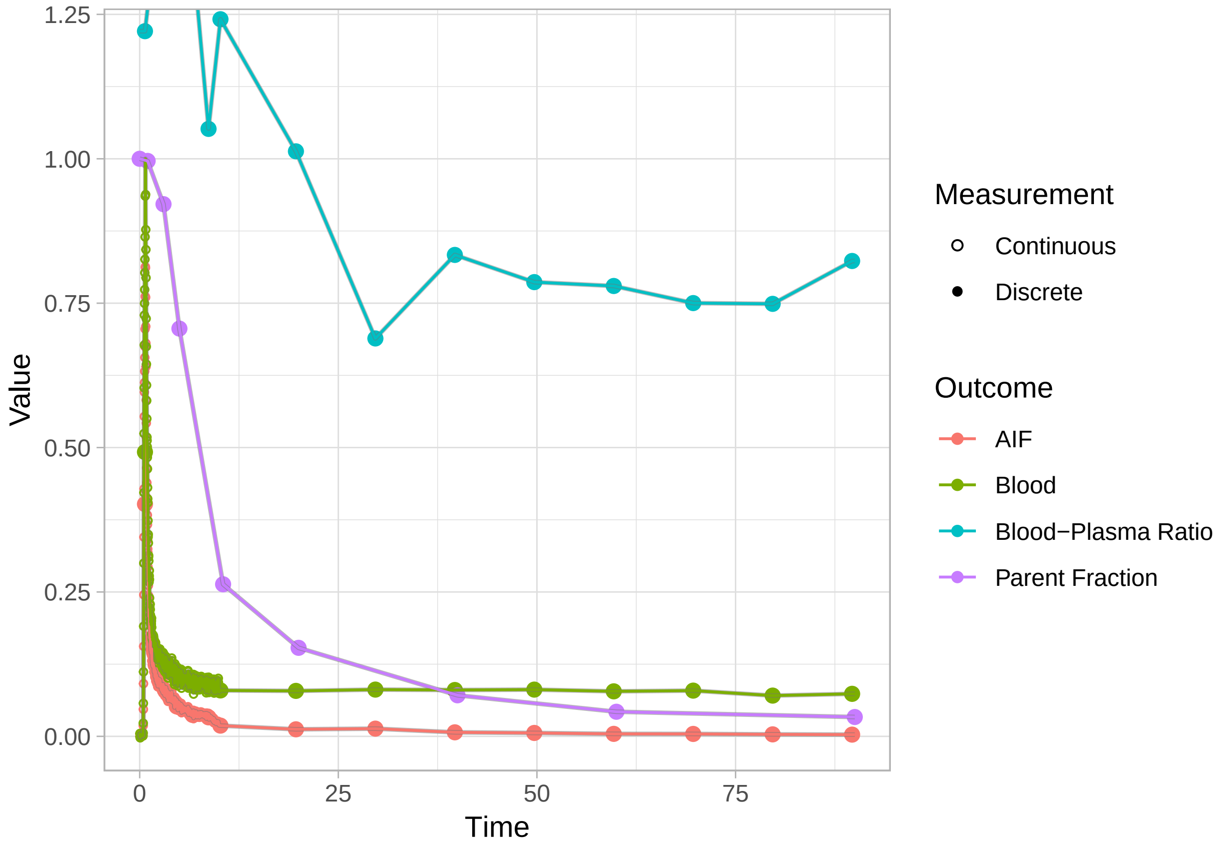


Let's first model the parent fraction, and then the blood-plasma ratio.

##### Modelling the parent fraction

Within *kinfitr*, there are a few different models for the parent fraction, which can be used at will, and compared easily using various model comparison metrics. For this exercise, I will just demonstrate fitting a model which was specifically designed for tracers within the same family as PBR28, namely the Guo modification of the Hill function. We'll fit it, and then add it to the *blooddata* object.

```
set.seed(123)

pbr28 <- pbr28 %>%
  group_by(PET) %>%
  mutate(pfdat = map(blooddata,
                 ~bd_getdata(.x, output = "parentFraction")),
         hillguomodel = map(pfdat,
                 ~metab_hillguo(.x$time, .x$parentFraction, multstart_iter = 100)),
         blooddata = map2(blooddata, hillguomodel,
                 ~bd_addfit(.x, .y, modeltype = "parentFraction"))) %>%
  select(-pfdat, -hillguomodel)
```

Now, we can check the *blooddata* plots again..

```
plot(pbr28$blooddata[[1]])
```

```
## Warning: Removed 1 rows containing missing values (geom_path).

## Warning: Removed 1 rows containing missing values (geom_path).
```

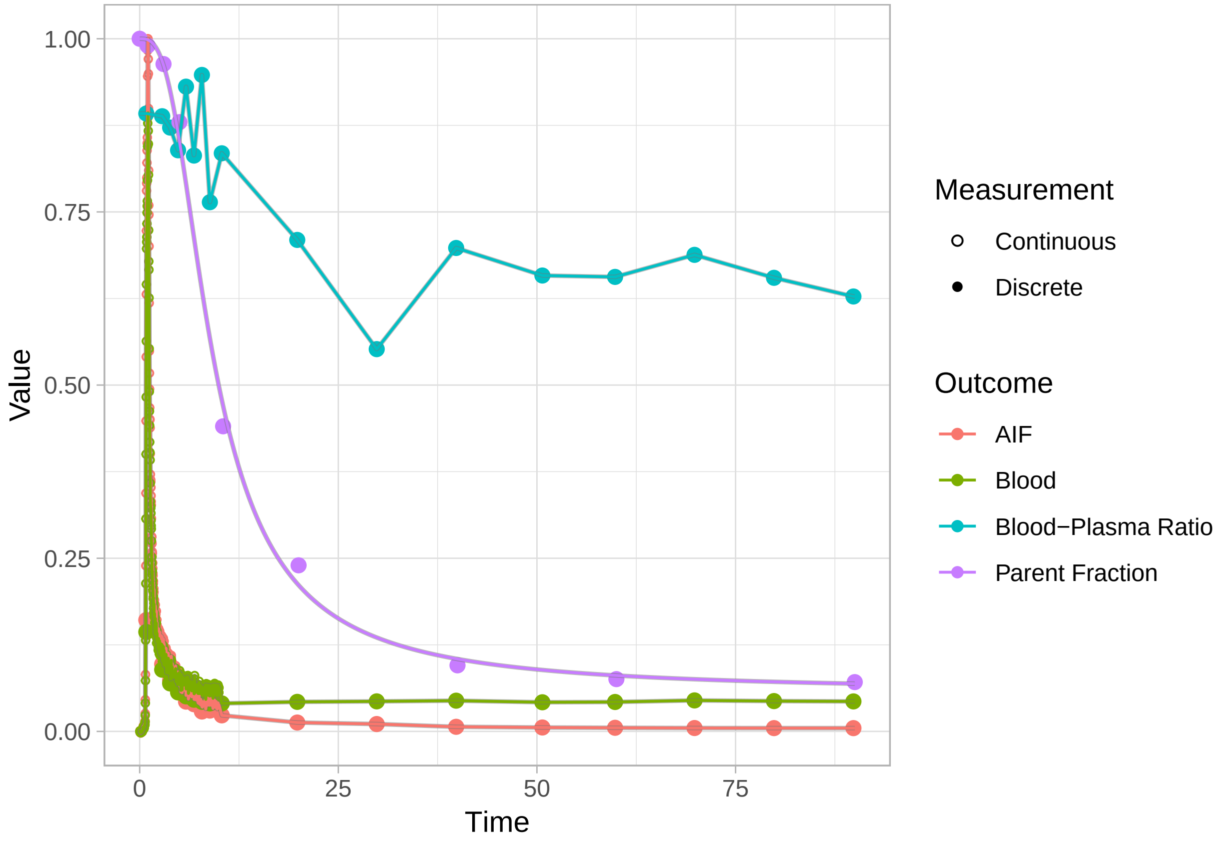


```
plot(pbr28$blooddata[[2]])
```

```
## Warning: Removed 1 rows containing missing values (geom_path).

## Warning: Removed 1 rows containing missing values (geom_path).
```

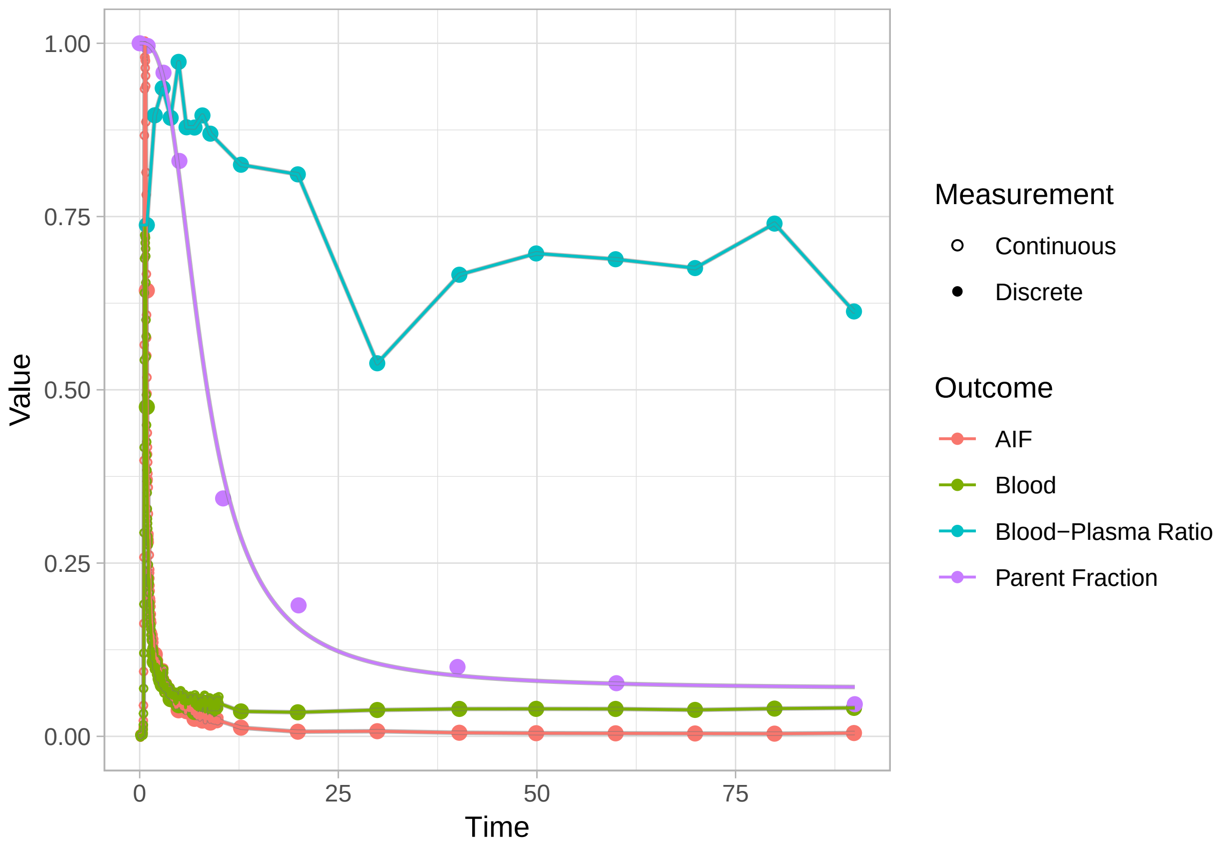


```r
plot(pbr28$blooddata[[3]])
```

```
## Warning: Removed 1 rows containing missing values (geom_path).

## Warning: Removed 1 rows containing missing values (geom_path).
```

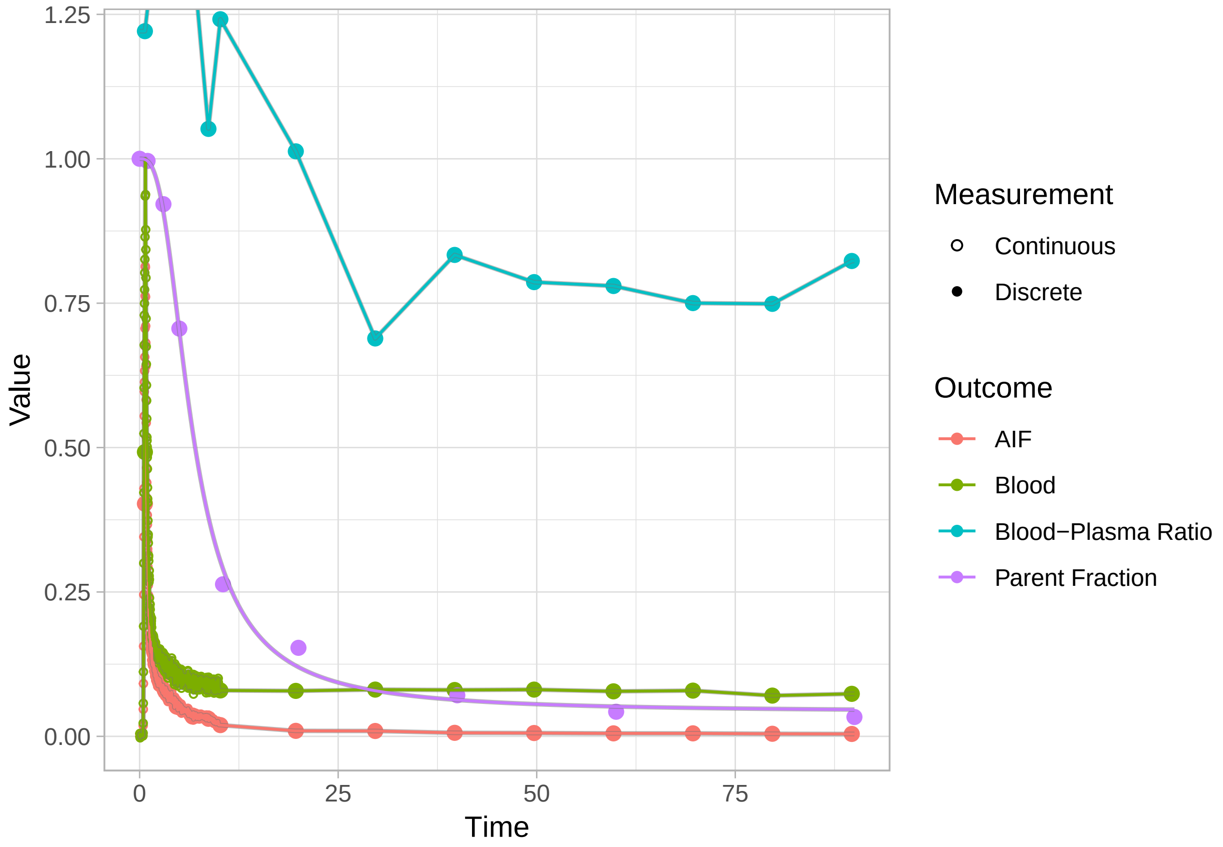


Now we see a much smoother fit to the parent fraction.

##### Modelling the blood-plasma ratio

The blood-plasma ratio doesn't seem to follow any particular function here, and there are no in-built methods for modelling this in *kinfitr*. However, kinfitr has been designed to make it easy to incorporate *ad-hoc* models which use the rest of the power of R for blood modelling. In this case, we can use a Generalised Additive Model, i.e. a smooth function, to model the blood-plasma ratio.

```
pbr28 <- pbr28 %>%
  group_by(PET) %>%
  mutate(bprdat = map(blooddata,
              ~bd_getdata(.x, output = "BPR")),
         bprmodel = map(bprdat,
              ~gam(bpr ~ s(time), data=.x)),
         blooddata = map2(blooddata, bprmodel,
              ~bd_addfit(.x, .y, modeltype = "BPR"))) %>%
  select(-bprdat, -bprmodel)
```

And let's check how the plots look:

```
plot(pbr28$blooddata[[1]])
```

```
## Warning: Removed 1 rows containing missing values (geom_path).

## Warning: Removed 1 rows containing missing values (geom_path).
```

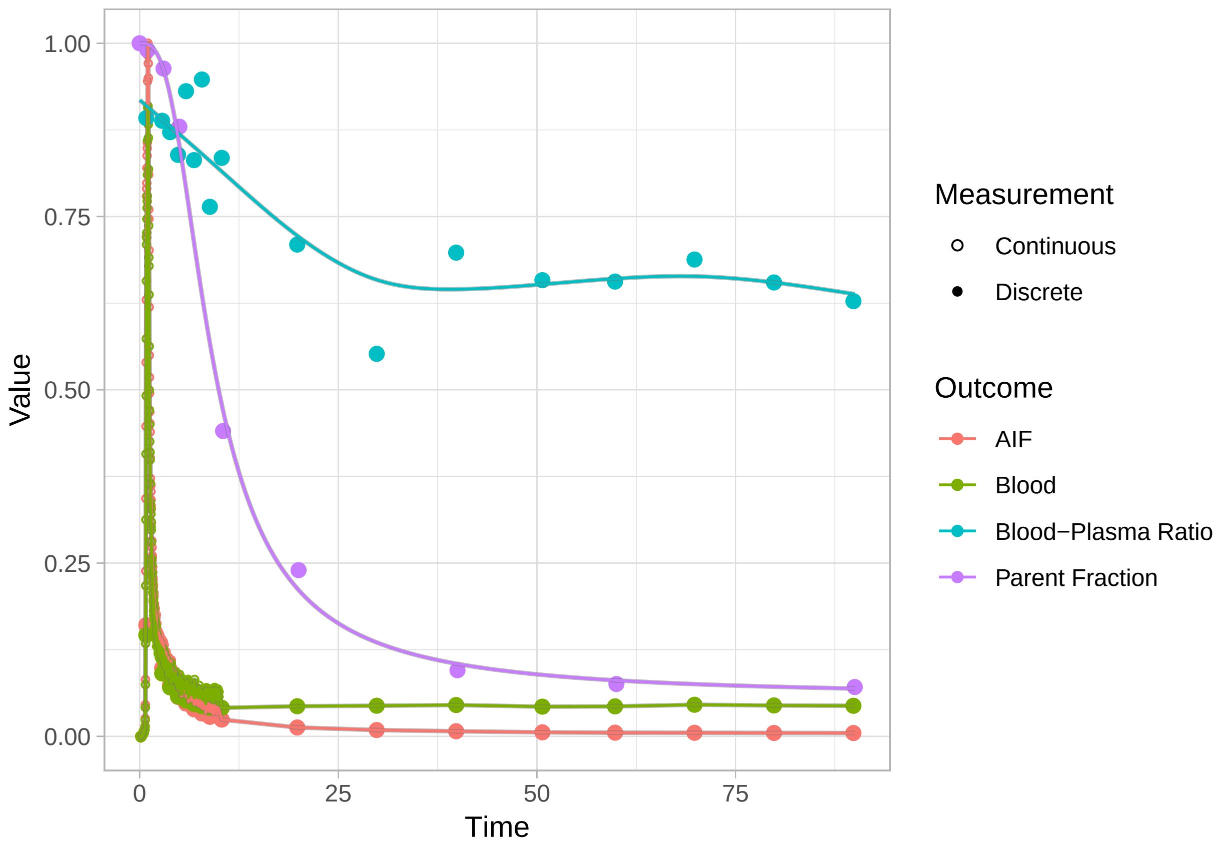


```
plot(pbr28$blooddata[[2]])
```

```
## Warning: Removed 1 rows containing missing values (geom_path).

## Warning: Removed 1 rows containing missing values (geom_path).
```

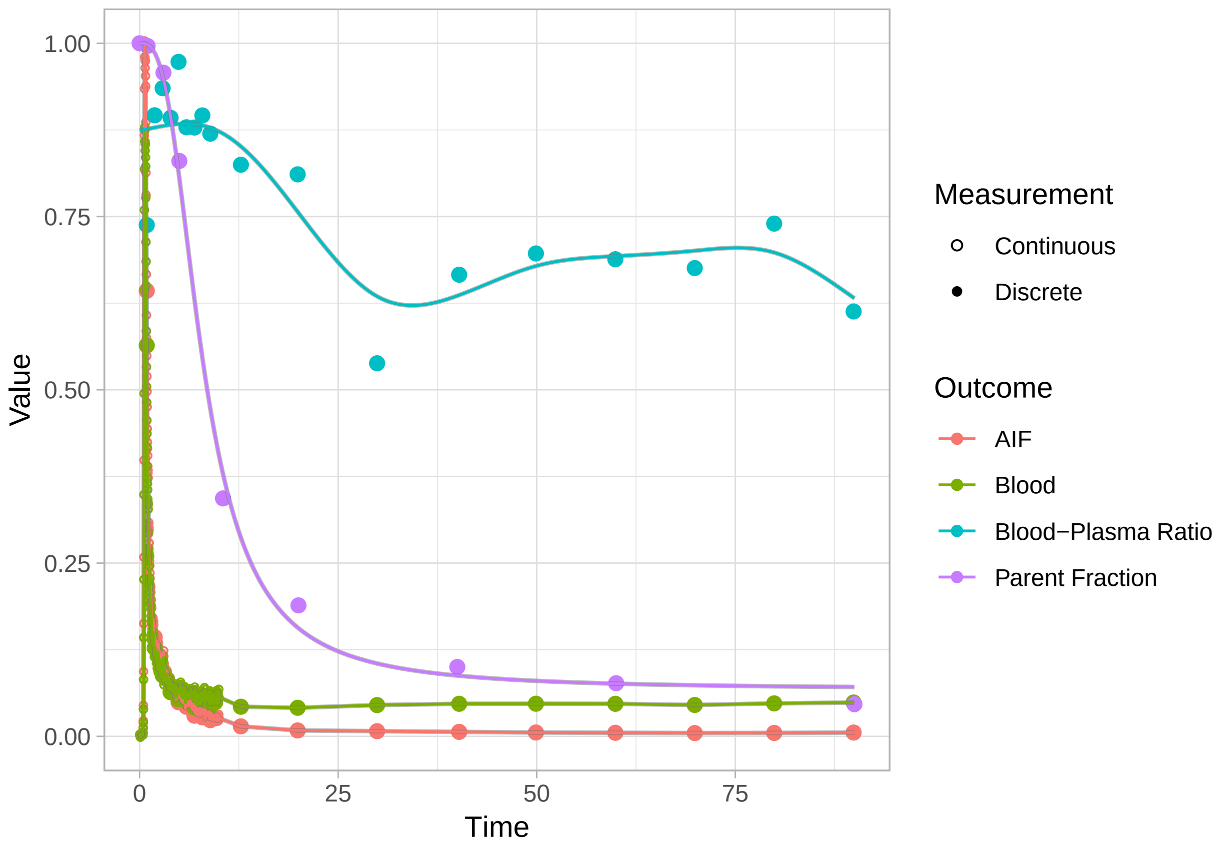


```r
plot(pbr28$blooddata[[3]])
```

```
## Warning: Removed 1 rows containing missing values (geom_path).

## Warning: Removed 1 rows containing missing values (geom_path).
```

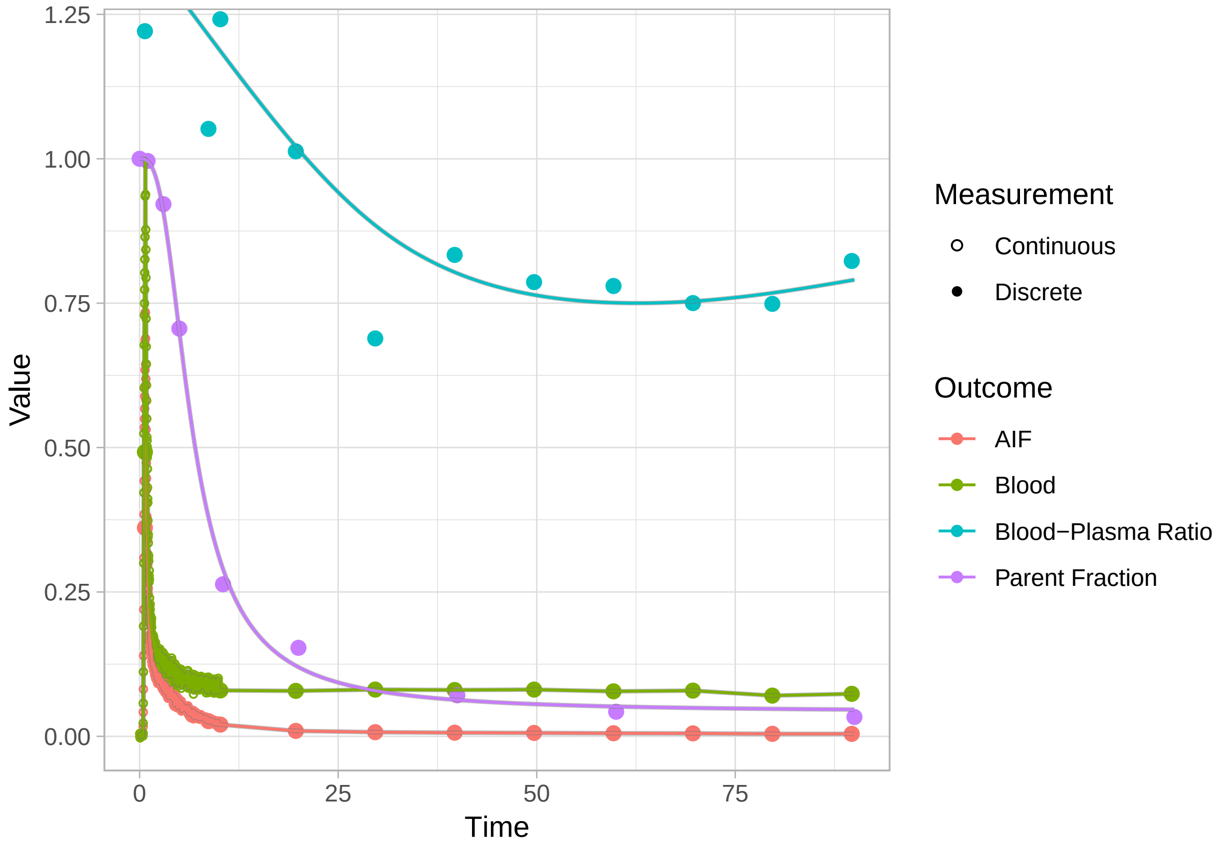


Clearly the fits are not fantastic, but they are certainly better than nothing.

##### Interpolating the blood data for kinetic modelling

Now, we want to extract the interpolated functions, in the same time units, for the purpose of using this in the kinetic models. This means, in *kinfitr*, extracting an *input* object from a *blooddata* object.

```
pbr28 <- pbr28 %>%
  group_by(PET) %>%
  mutate(input = map(blooddata, bd_getdata))
```

Done! Now we can get to modelling.

#### Preparations for Modelling the TACs

#### Modelling the Delay

The first modelling of the TAC which is necessary is to estimate the delay between the blood samples and the TAC. We can do this using one of the tissue compartmental models, with an additional parameter representing the delay.

```
pbr28 <- pbr28 %>%
  group_by(Subjname) %>%
  mutate(delayFit = map2(tacs, input,
                         ~twotcm(t_tac = .x$Times/60, # sec to min
                                 tac = .x$WB,
                                 input = .y,
                                 weights = .x$Weights,
```

```
                                vB=0.05)))
```

```
## Warning in twotcm(t_tac = .x$Times/60, tac = .x$WB, input = .y, weights = .x$Weights, : Fitted parame
## either modifying the upper and lower limit boundaries, or else using
## multstart when fitting the model (see the function documentation).

## Warning in twotcm(t_tac = .x$Times/60, tac = .x$WB, input = .y, weights = .x$Weights, : Fitted parame
## either modifying the upper and lower limit boundaries, or else using
## multstart when fitting the model (see the function documentation).
```

Now let's check them, because delay fitting can go slightly awry.

```
walk2(pbr28$delayFit, pbr28$PET ,
      ~print(plot_inptac_fit(.x) + ggtitle(.y)))
```

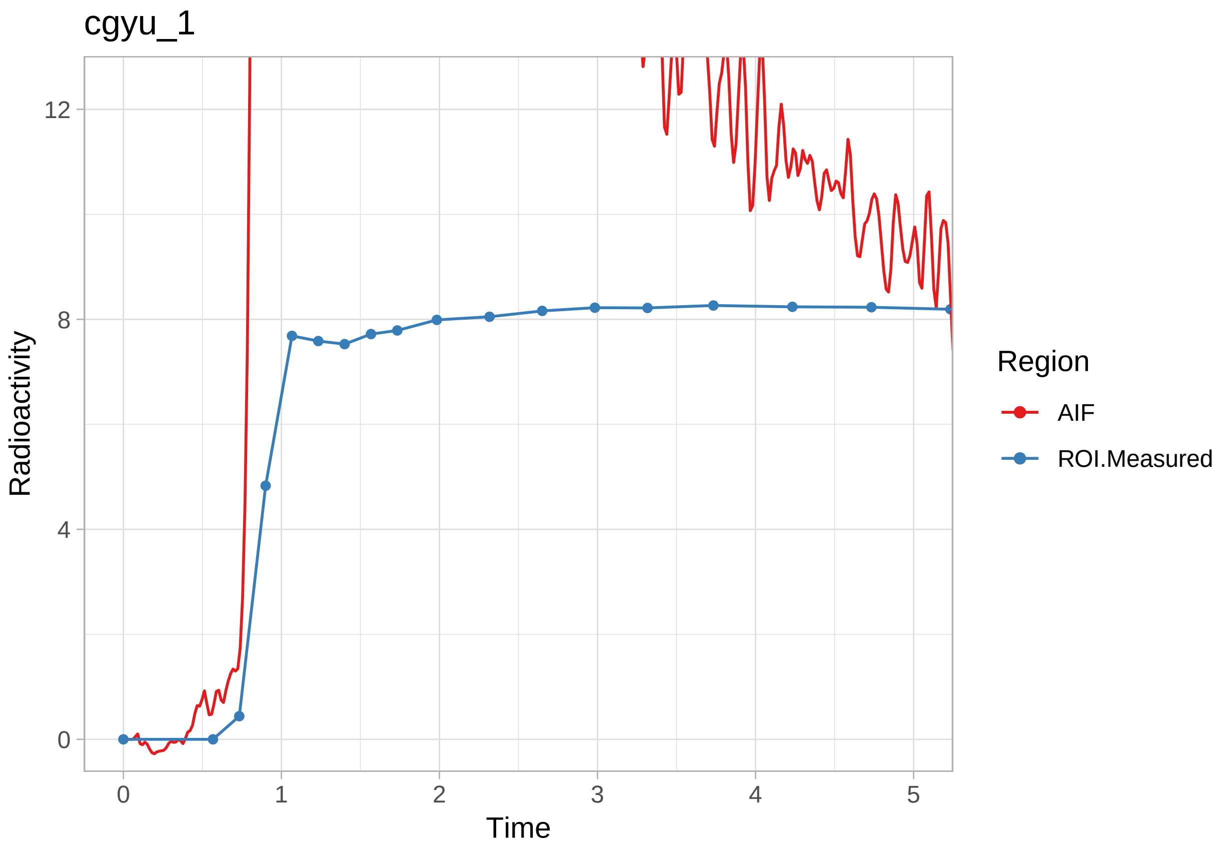

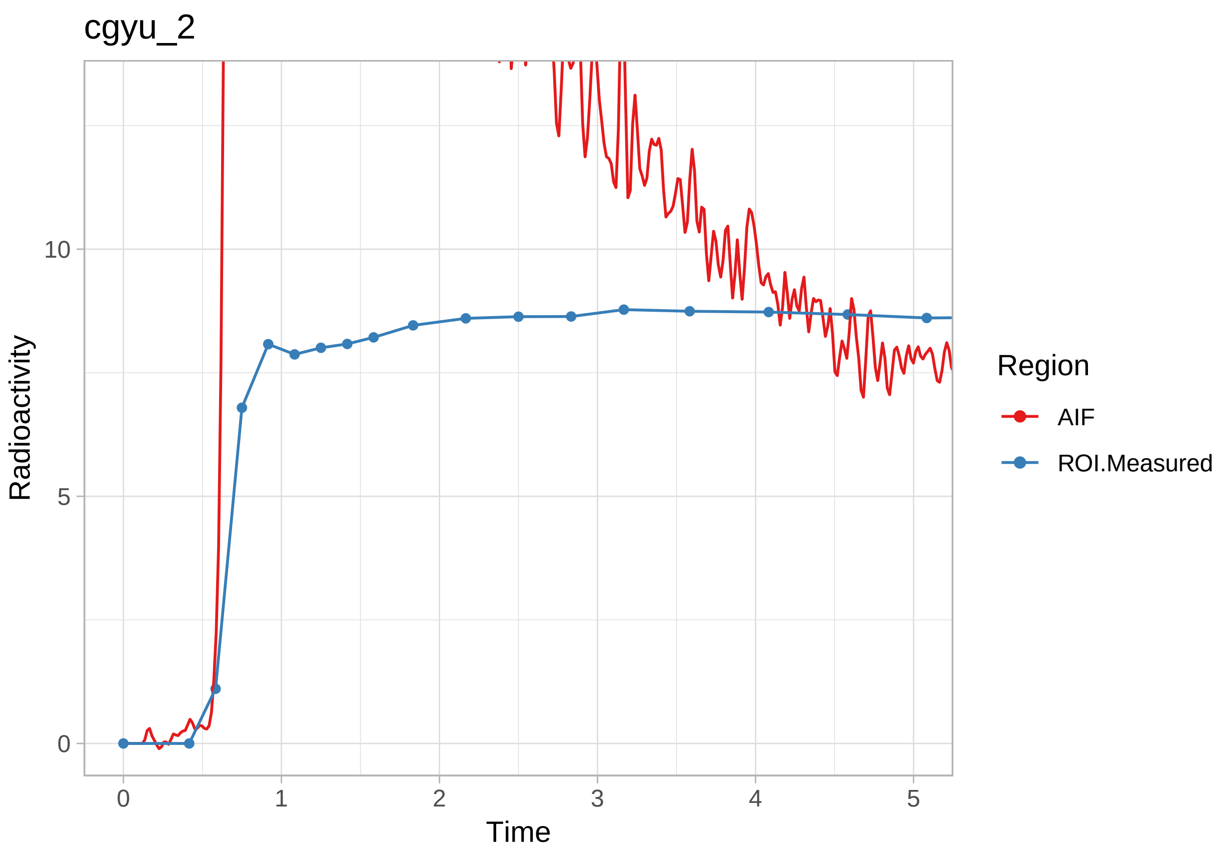
cgyu_2
Radioactivity
Time
0
5
10
0
1
2
3
4
5
Region
AIF
ROI.Measured

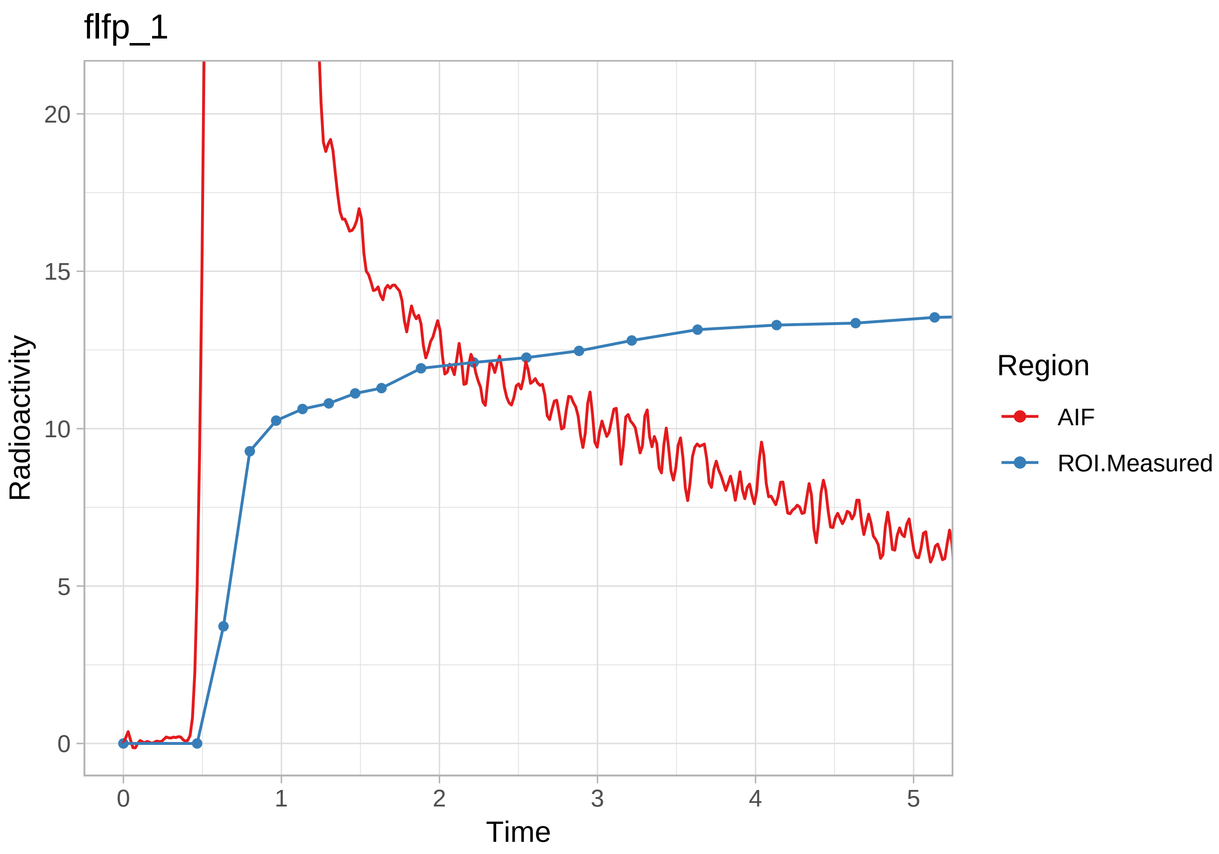
flfp_1
Radioactivity
Time
0
5
10
15
20
0
1
2
3
4
5
Region
AIF
ROI.Measured

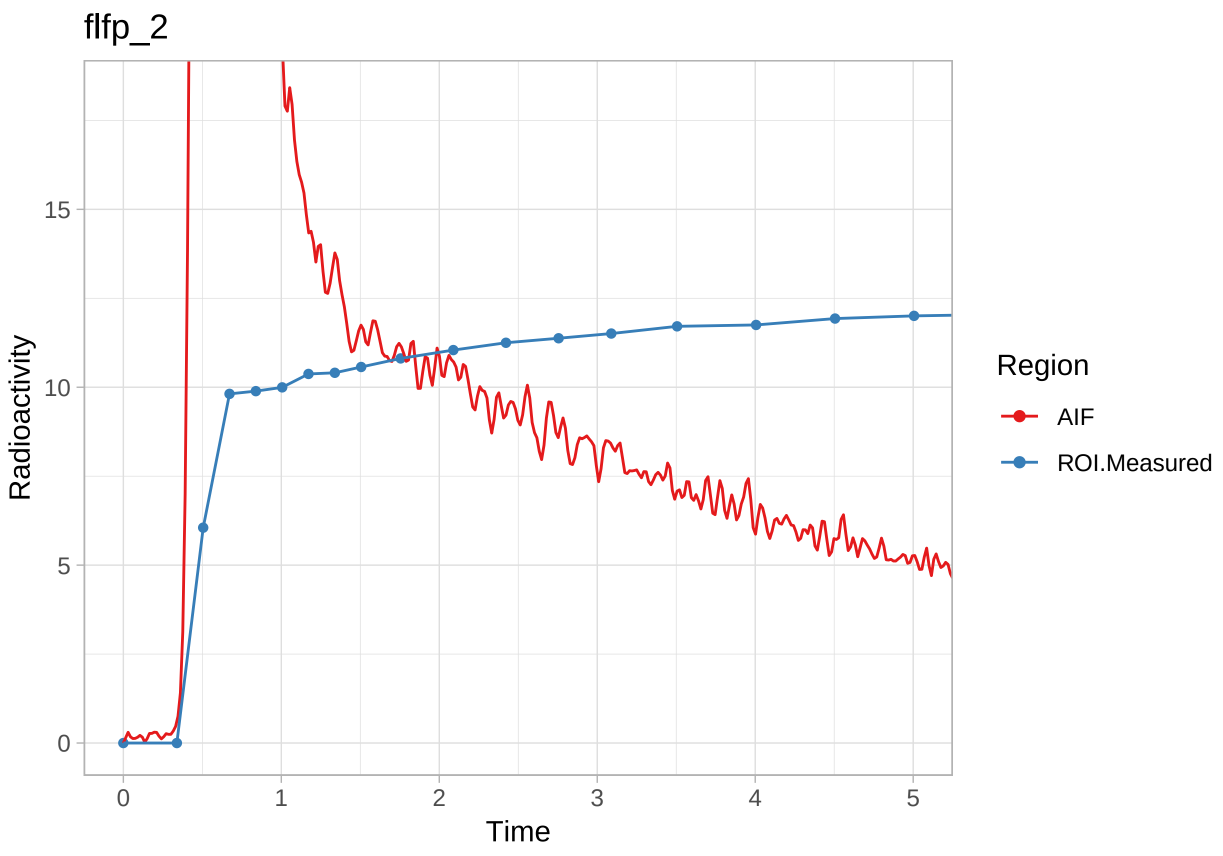
flfp_2
Radioactivity
0
5
10
15
0
1
2
3
4
5
Time
Region
AIF
ROI.Measured

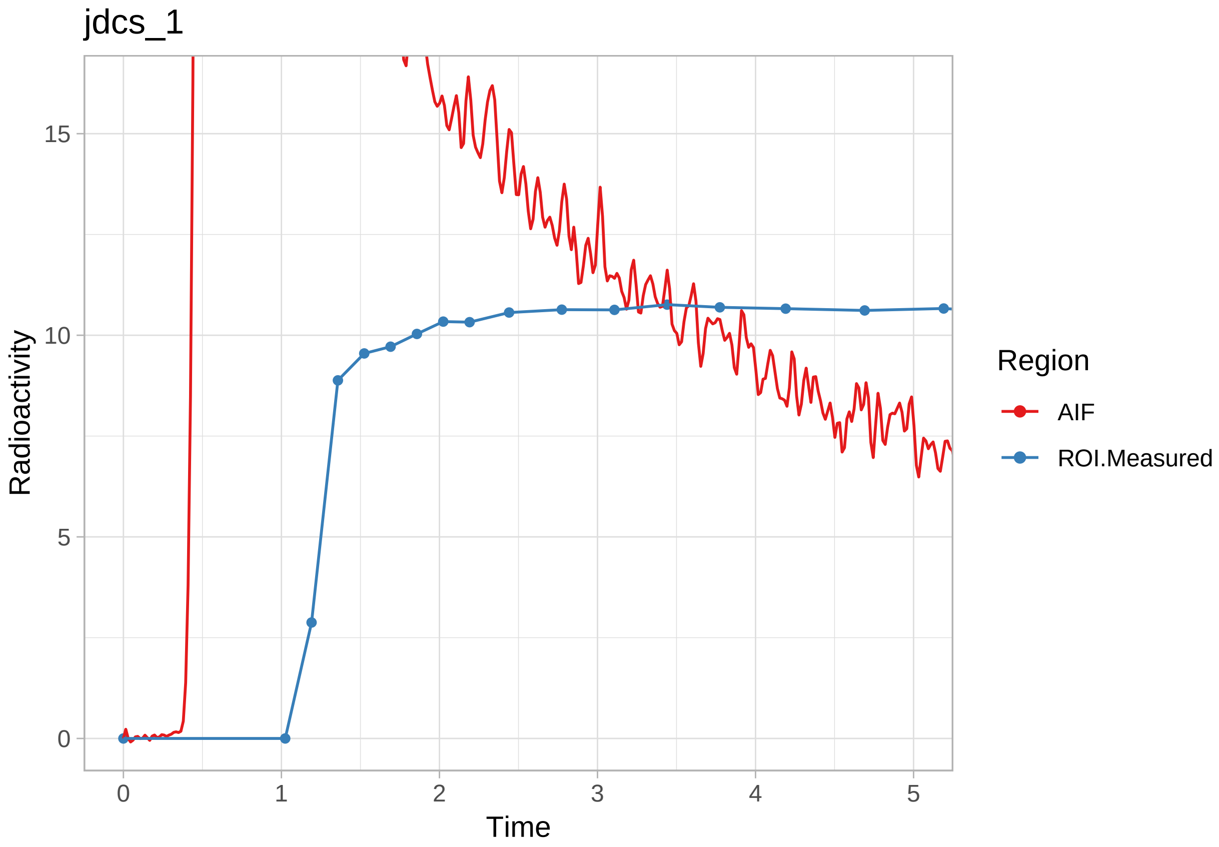
jdcs_1
Radioactivity
Time
Region
AIF
ROI.Measured
0
5
10
15
0
1
2
3
4
5

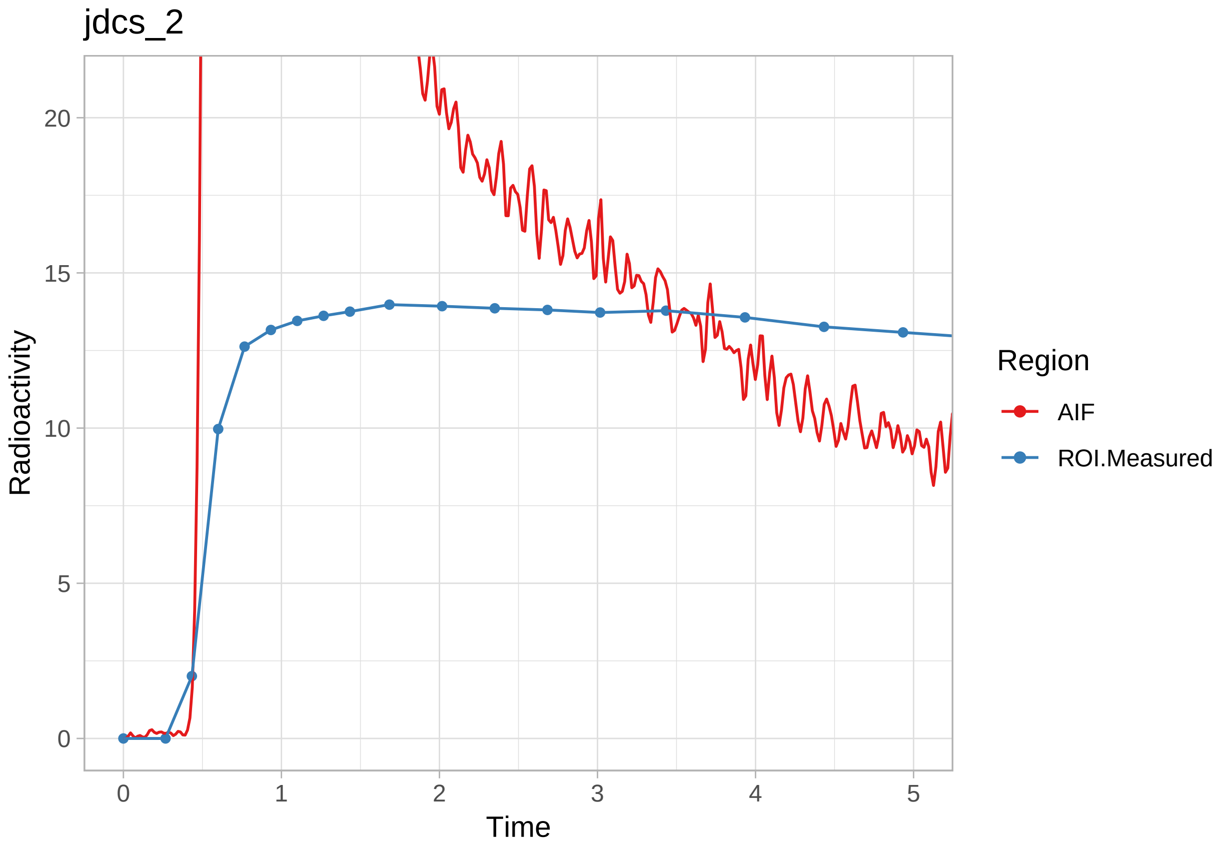
jdcs_2
Radioactivity
Time
Region
AIF
ROI.Measured
0
5
10
15
20
0
1
2
3
4
5

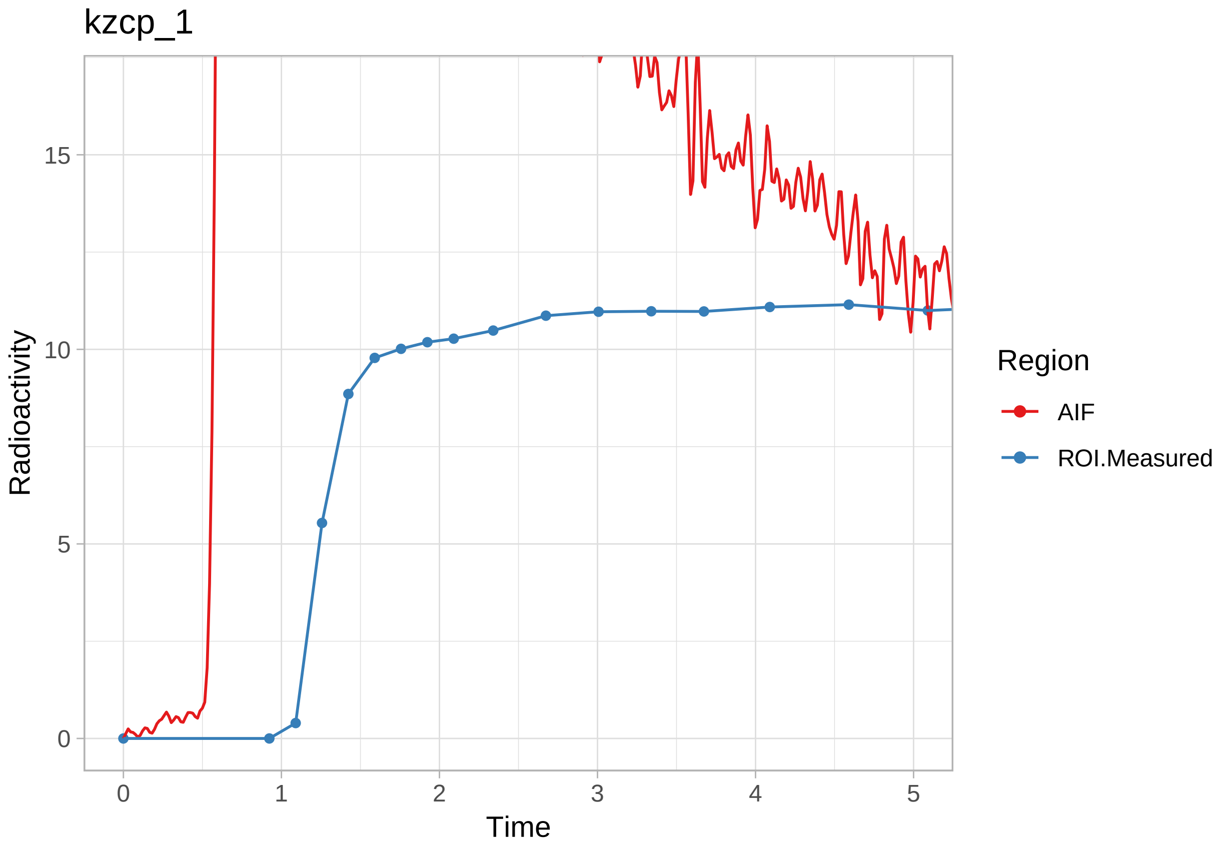

kzcp_1
Radioactivity
15
10
5
0
0
1
2
3
4
5
Time
Region
AIF
ROI.Measured

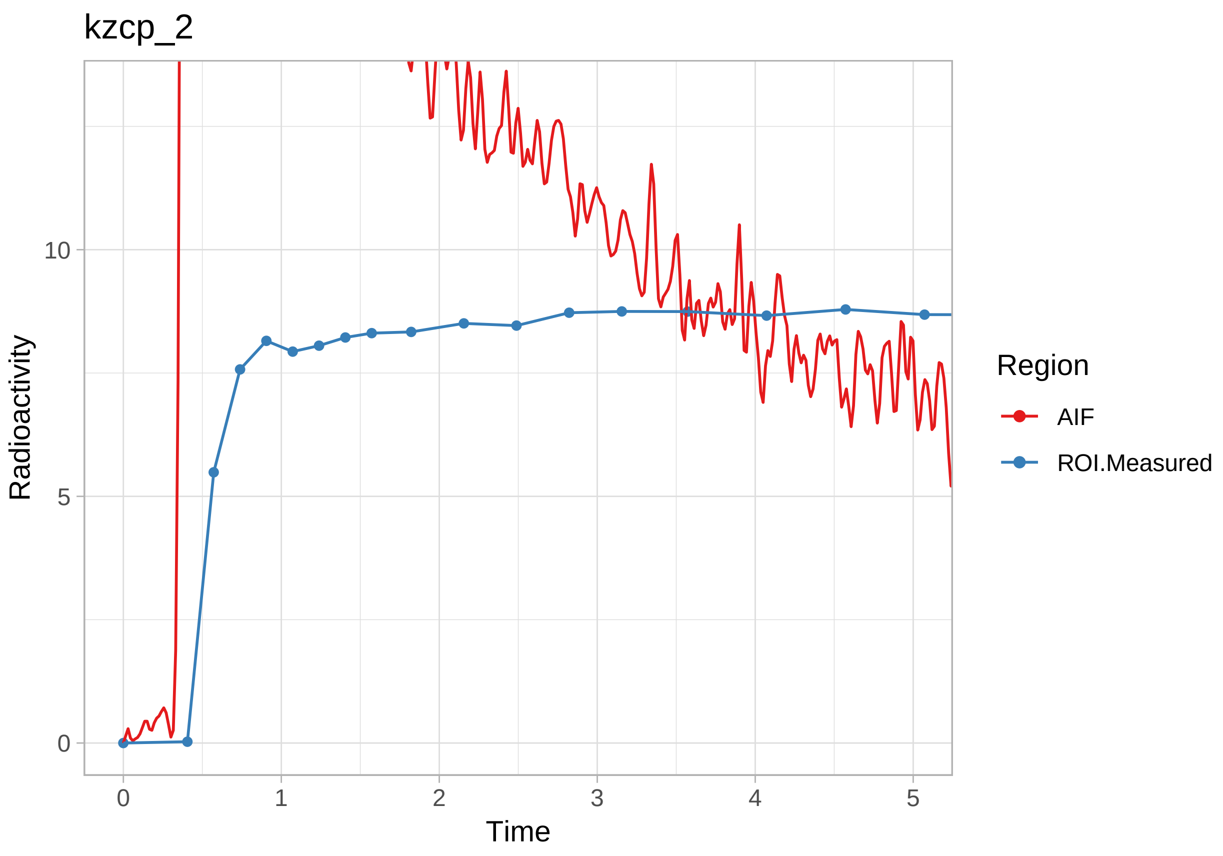
kzcp_2
Radioactivity
10
5
0
0
1
2
3
4
5
Time
Region
AIF
ROI.Measured

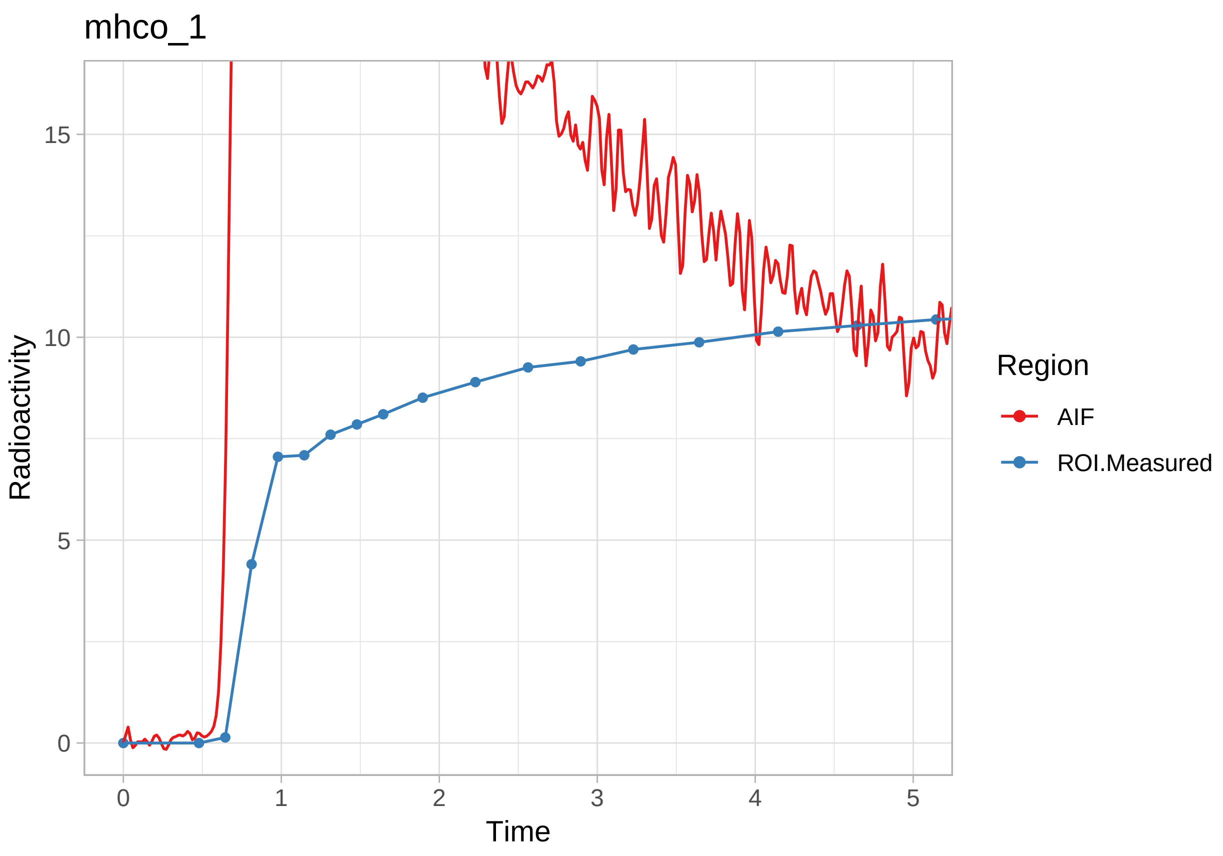
mhco_1
Radioactivity
0
5
10
15
0
1
2
3
4
5
Time
Region
AIF
ROI.Measured

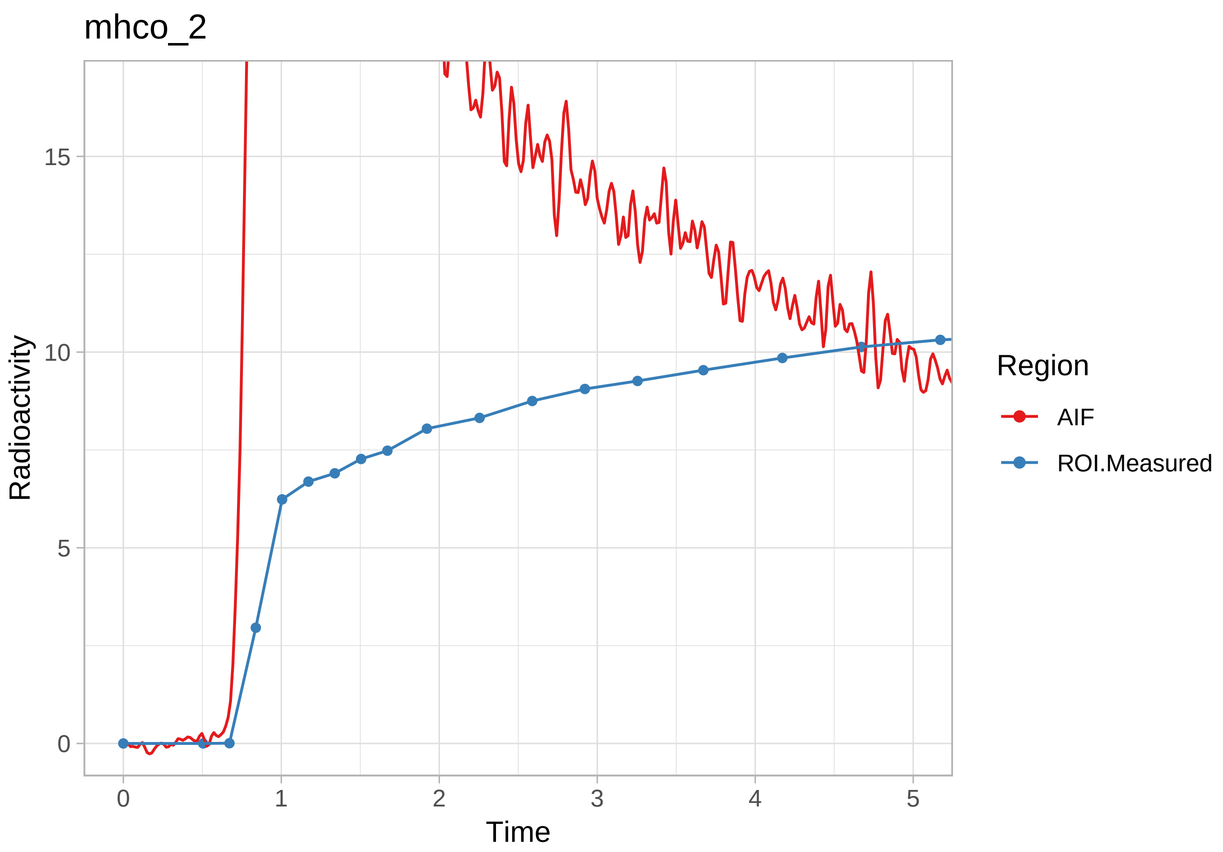
mhco_2
Radioactivity
15
10
5
0
0
1
2
3
4
5
Time
Region
AIF
ROI.Measured

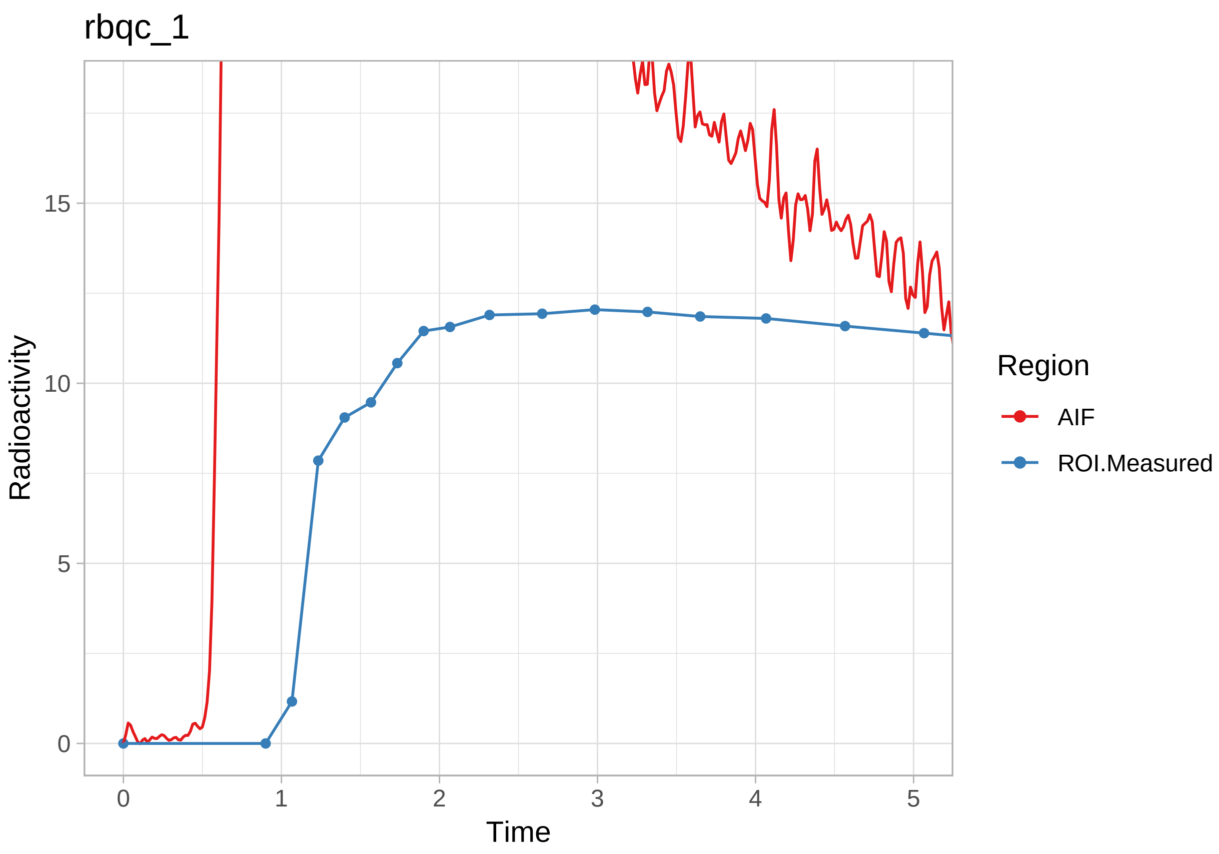

rbqc_1
Radioactivity
15
10
5
0
0
1
2
3
4
5
Time
Region
AIF
ROI.Measured

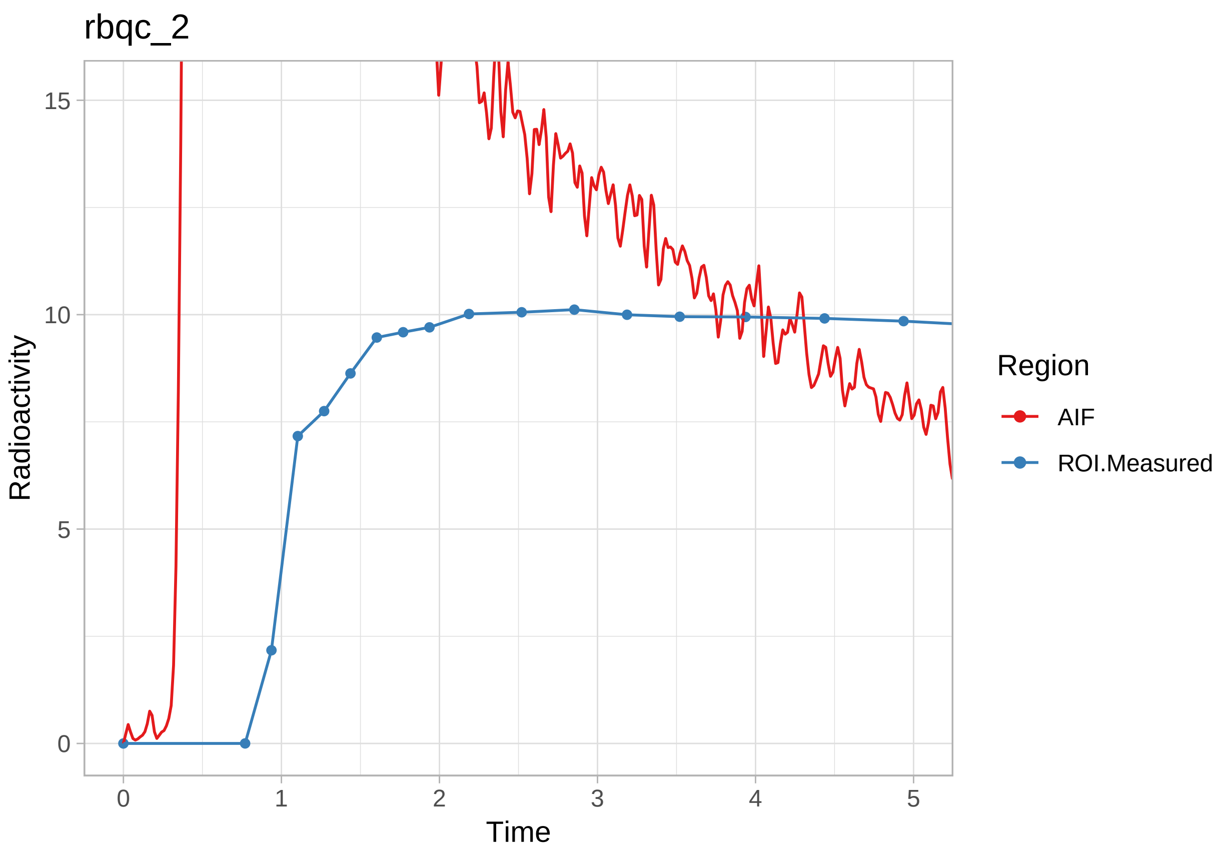
rbqc_2
Radioactivity
Time
Region
AIF
ROI.Measured
15
10
5
0
0
1
2
3
4
5

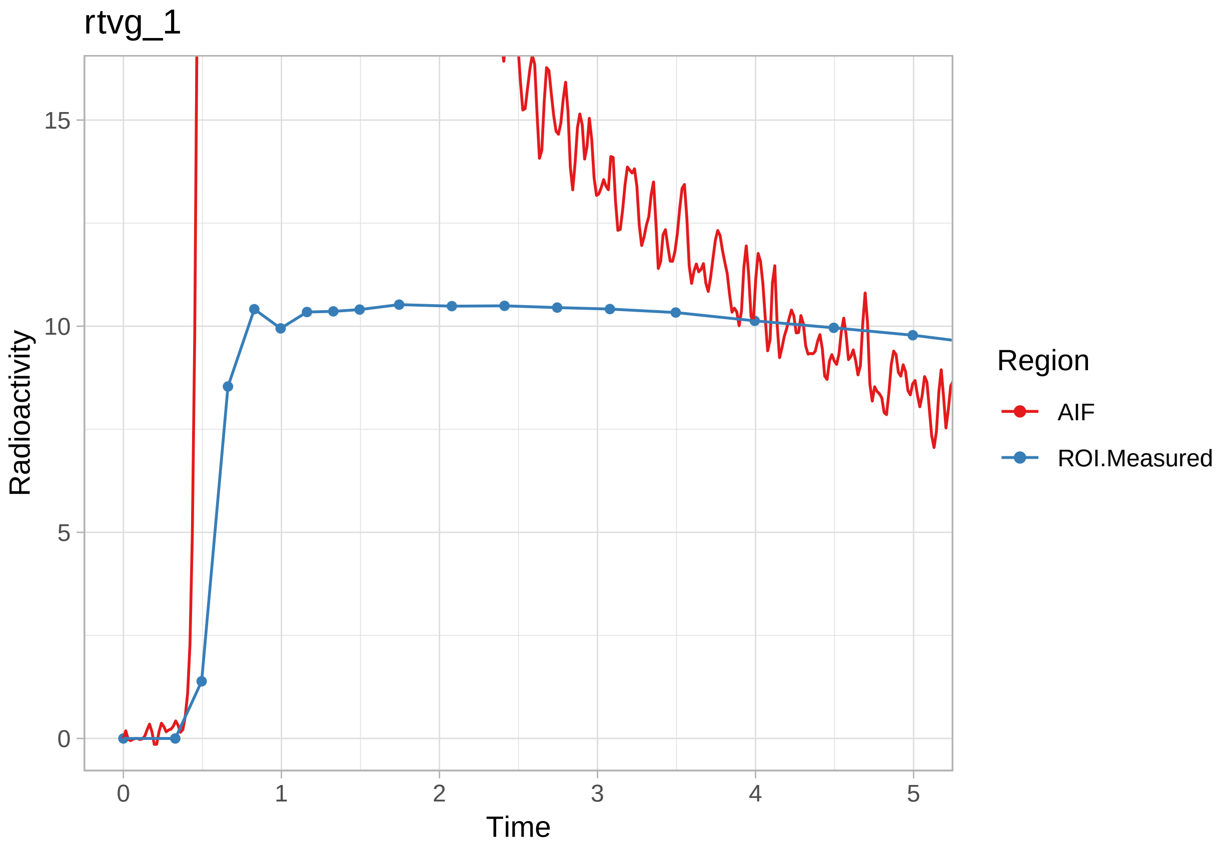
rtvg_1
Radioactivity
15
10
5
0
0
1
2
3
4
5
Time
Region
AIF
ROI.Measured

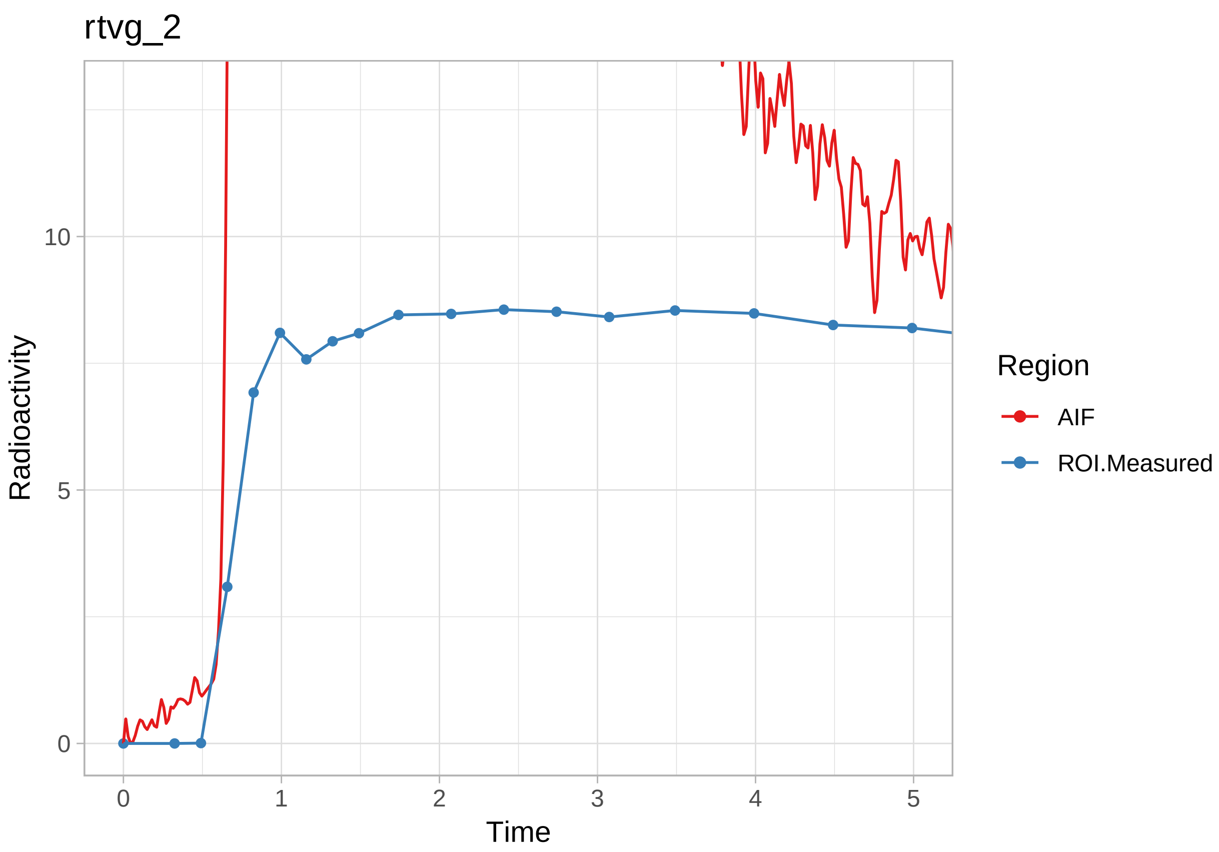
rtvg_2
Radioactivity
10
5
0
0
1
2
3
4
5
Time
Region
AIF
ROI.Measured

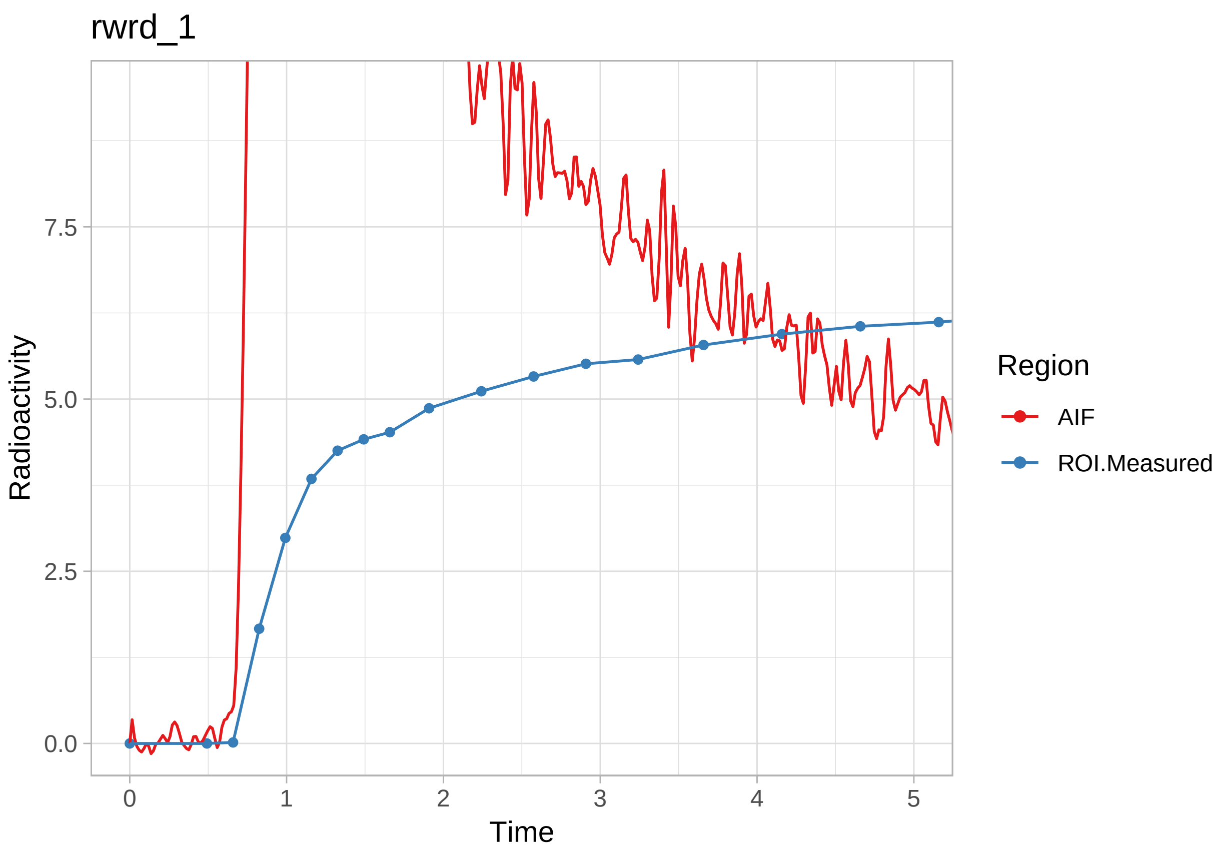
rwrd_1
Radioactivity
0.0
2.5
5.0
7.5
0
1
2
3
4
5
Time
Region
AIF
ROI.Measured

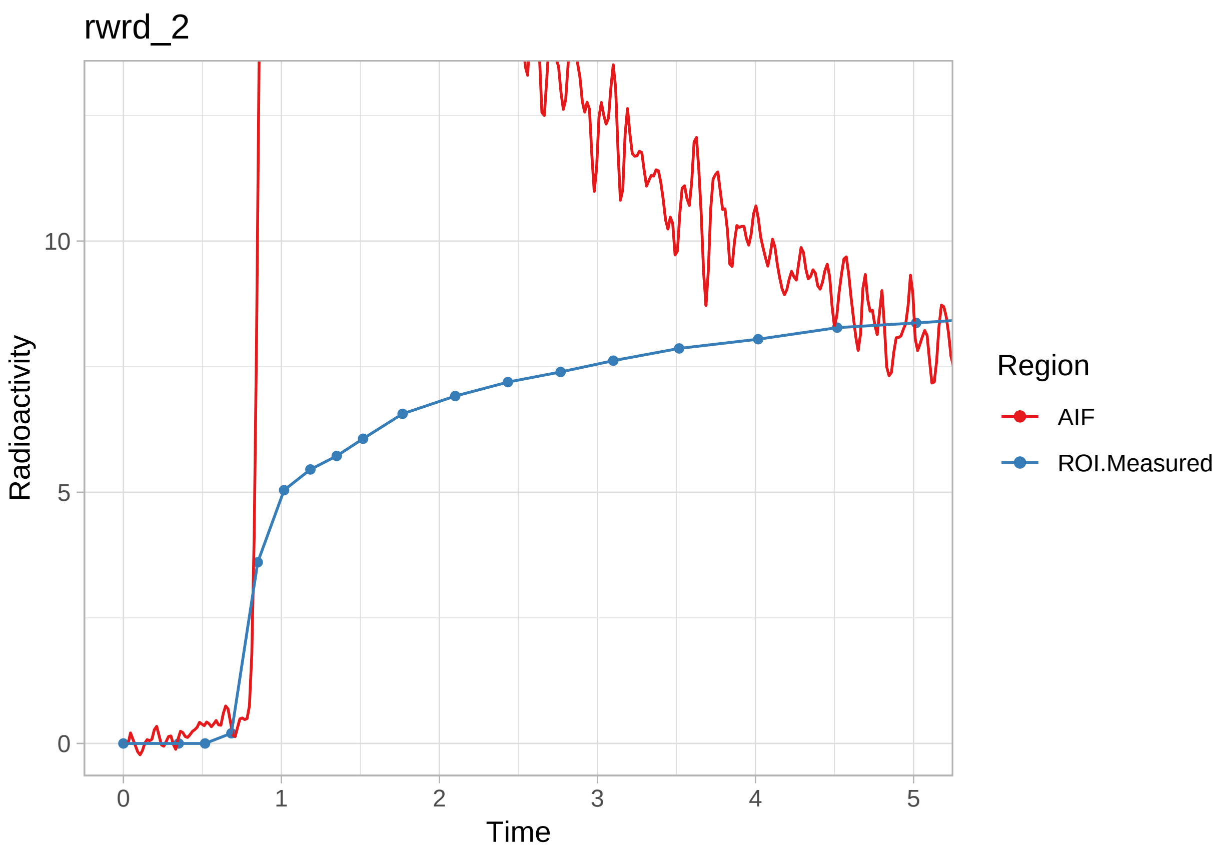
rwrd_2
Radioactivity
10
5
0
0
1
2
3
4
5
Time
Region
AIF
ROI.Measured

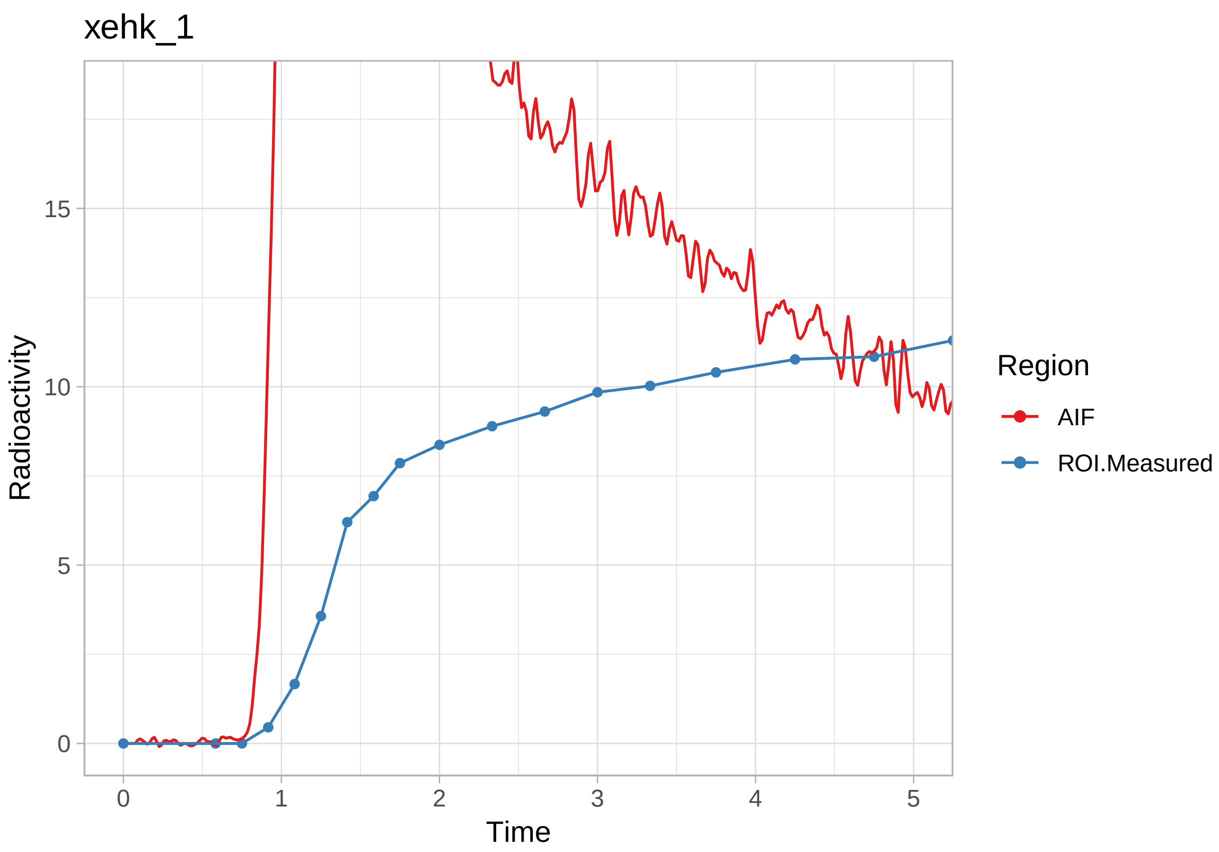
xehk_1
Radioactivity
Time
0
5
10
15
0
1
2
3
4
5
Region
AIF
ROI.Measured

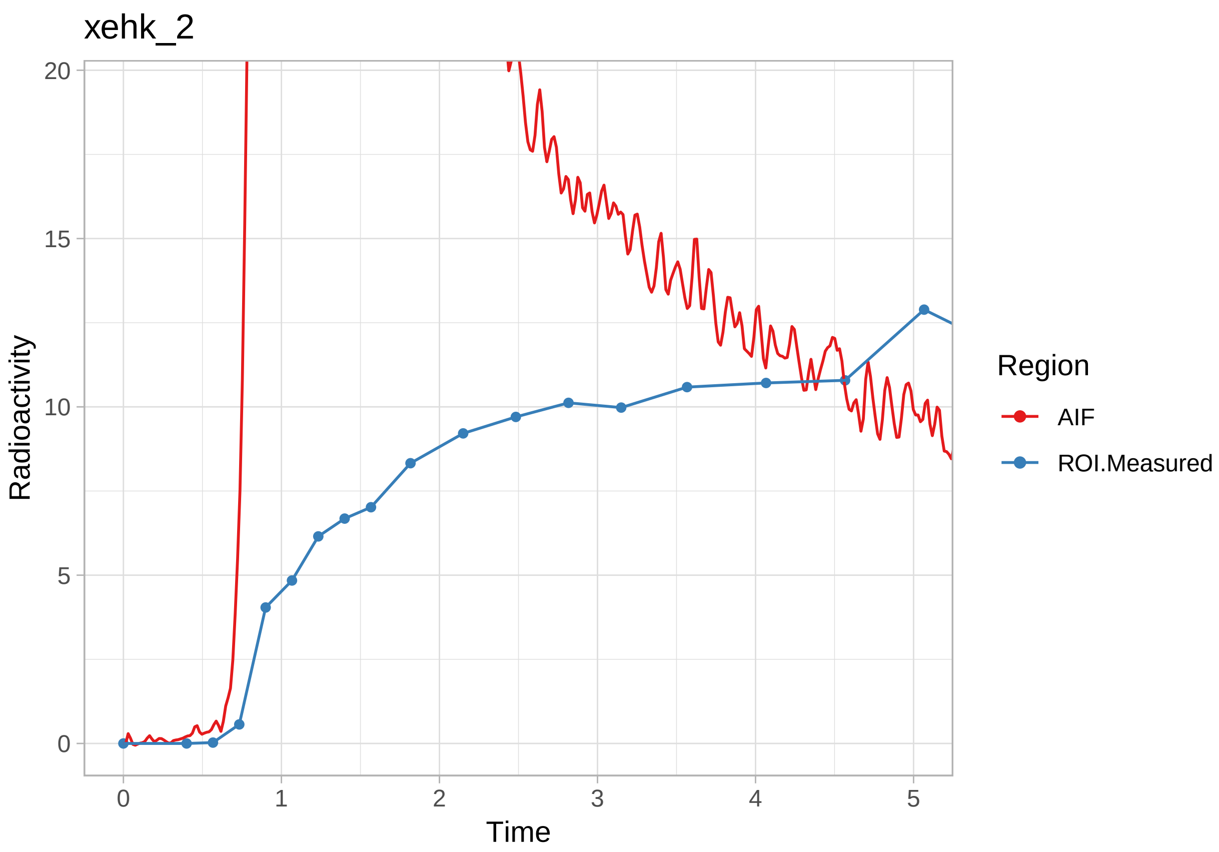

xehk_2
Radioactivity
20
15
10
5
0
0
1
2
3
4
5
Time
Region
AIF
ROI.Measured

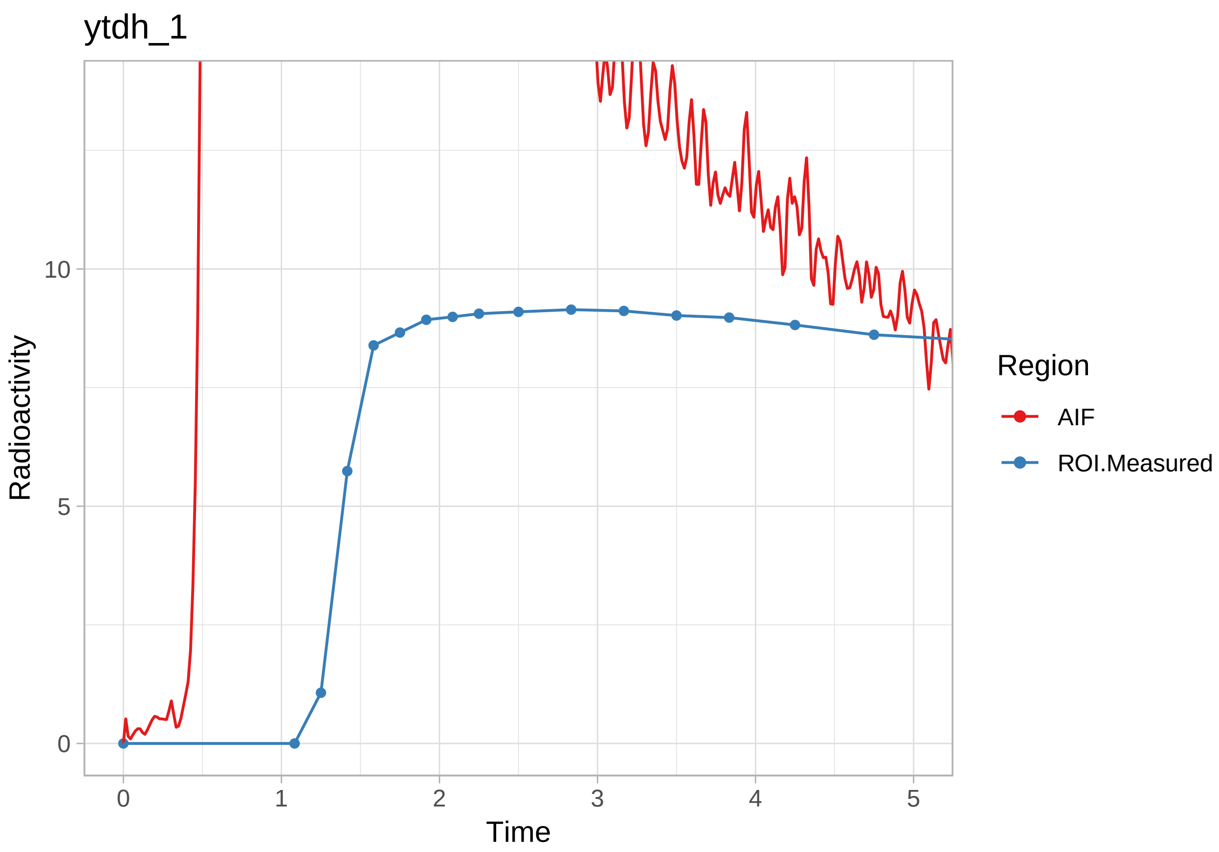
ytdh_1
Radioactivity
10
5
0
0
1
2
3
4
5
Time
Region
AIF
ROI.Measured

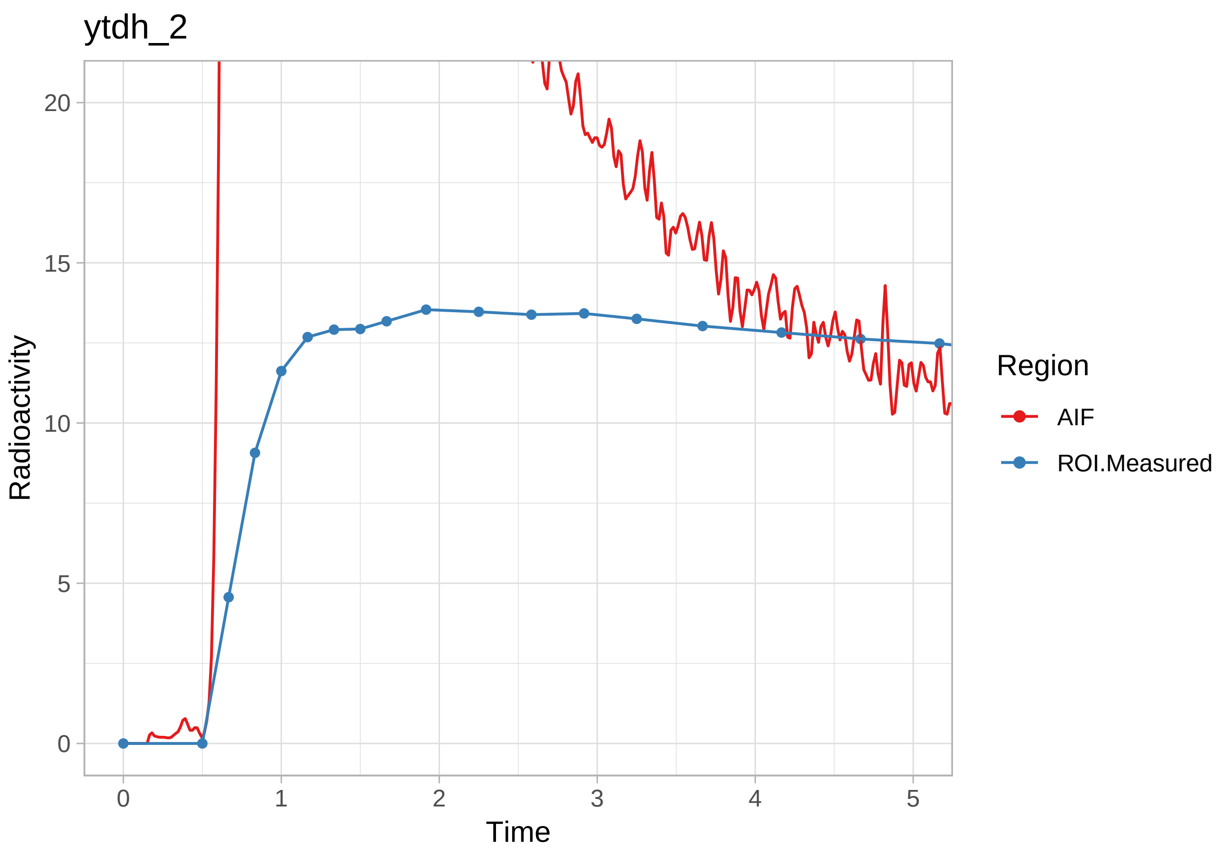


OK, those look ok! Now we'll extract the delay values and use those in future models.

```
pbr28 <- pbr28 %>%
  group_by(Subjname) %>%
  mutate(inpshift = map_dbl(delayFit, c("par", "inpshift")))
```

##### Estimating t*

For the linear models, we will need to decide on t* values again. Instead of doing it again in this tutorial, we will just use 10 frames.

#### Model Fitting

Now that we have interpolated the blood, modelled the delay and selected t* values, we are ready to start modelling the TACs. Let's first select three TACs per individual.

```
pbr28 <- pbr28 %>%
  group_by(PET) %>%
  mutate(tacs = map(tacs, ~select(.x, Times, Weights, FC, STR, CBL))) %>%
  select(PET, tacs, input, inpshift)
```

Now we have to rearrange the data as before, but now we also have the input data. Let's first separate the data, rearrange, and then join them again.

```
pbr28_input <- select(pbr28, PET, input)

pbr28_tacs <- select(pbr28, PET, tacs, inpshift)
```

```
pbr28_long <- pbr28_tacs %>%
  unnest() %>%
  gather(key = Region, value = TAC, -Times, -Weights, -inpshift, -PET) %>%
  group_by(PET, inpshift, Region) %>%
  nest(.key = "tacdata") %>%
  full_join(pbr28_input)
```

```
## Joining, by = "PET"
```

And now let's see what it looks like now.

```
pbr28_long
```

```
## # A tibble: 60 x 5
##    PET    inpshift Region tacdata           input
##    <chr>     <dbl> <chr>  <list>            <list>
##  1 cgyu_1  0.0613  FC     <tibble [38 x 3]> <tibble [6,000 x 5]>
##  2 cgyu_2  0.130   FC     <tibble [38 x 3]> <tibble [6,000 x 5]>
##  3 flfp_1 -0.0666  FC     <tibble [38 x 3]> <tibble [6,000 x 5]>
##  4 flfp_2 -0.00544 FC     <tibble [38 x 3]> <tibble [6,000 x 5]>
##  5 jdcs_1 -0.457   FC     <tibble [38 x 3]> <tibble [6,000 x 5]>
##  6 jdcs_2  0.0202  FC     <tibble [38 x 3]> <tibble [6,000 x 5]>
##  7 kzcp_1 -0.257   FC     <tibble [38 x 3]> <tibble [6,000 x 5]>
##  8 kzcp_2 -0.0221  FC     <tibble [38 x 3]> <tibble [6,000 x 5]>
##  9 mhco_1 -0.262   FC     <tibble [38 x 3]> <tibble [6,000 x 5]>
## 10 mhco_2 -0.205   FC     <tibble [38 x 3]> <tibble [6,000 x 5]>
## # ... with 50 more rows
```

Now we can proceed to fitting models to all the regions. Where we used the *purrr::map* and *purrr::map2* functions before, we must now use *purrr::pmap*. This will require us to write a few small functions first. We'll also just skip returning the whole model fits, and just return the $V_T$ values.

```
fit_1tcm <- function(tacdata, input, inpshift) {
  onetcm(t_tac = tacdata$Times/60, tac = tacdata$TAC,
         input = input, weights = tacdata$Weights,
         inpshift = inpshift)$par$Vt
}

fit_2tcm <- function(tacdata, input, inpshift) {
  twotcm(t_tac = tacdata$Times/60, tac = tacdata$TAC,
         input = input, weights = tacdata$Weights,
         inpshift = inpshift)$par$Vt
}

fit_2tcm1k <- function(tacdata, input, inpshift) {
  twotcm1k(t_tac = tacdata$Times/60, tac = tacdata$TAC,
         input = input, weights = tacdata$Weights,
         inpshift = inpshift, vB = 0.05)$par$Vt
}

fit_Logan <- function(tacdata, input, inpshift) {
  Loganplot(t_tac = tacdata$Times/60, tac = tacdata$TAC,
         input = input, weights = tacdata$Weights,
         inpshift = inpshift,
         tstarIncludedFrames = 10)$par$Vt
}
```

```
fit_mlLogan <- function(tacdata, input, inpshift) {
  mlLoganplot(t_tac = tacdata$Times/60, tac = tacdata$TAC,
         input = input, weights = tacdata$Weights,
         inpshift = inpshift,
         tstarIncludedFrames = 10)$par$Vt
}

fit_ma1 <- function(tacdata, input, inpshift) {
  ma1(t_tac = tacdata$Times/60, tac = tacdata$TAC,
         input = input, weights = tacdata$Weights,
         inpshift = inpshift,
         tstarIncludedFrames = 10)$par$Vt
}

fit_ma2 <- function(tacdata, input, inpshift) {
  ma2(t_tac = tacdata$Times/60, tac = tacdata$TAC,
         input = input, weights = tacdata$Weights,
         inpshift = inpshift)$par$Vt
}
```

And now let's fit the models. We'll also separate this one out for the purposes of seeing where the boundary limits are hit.

```
pbr28_long <- pbr28_long %>%
  group_by(PET, Region)

  # 1TCM
pbr28_long <- pbr28_long %>%
  mutate("1TCM" = pmap_dbl(list(tacdata, input, inpshift), fit_1tcm))
```

```
## Warning in onetcm(t_tac = tacdata$Times/60, tac = tacdata$TAC, input = input, : Fitted parameters ar
## either modifying the upper and lower limit boundaries, or else using
## multstart when fitting the model (see the function documentation).

## Warning in onetcm(t_tac = tacdata$Times/60, tac = tacdata$TAC, input = input, : Fitted parameters ar
## either modifying the upper and lower limit boundaries, or else using
## multstart when fitting the model (see the function documentation).

## Warning in onetcm(t_tac = tacdata$Times/60, tac = tacdata$TAC, input = input, : Fitted parameters ar
## either modifying the upper and lower limit boundaries, or else using
## multstart when fitting the model (see the function documentation).

## Warning in onetcm(t_tac = tacdata$Times/60, tac = tacdata$TAC, input = input, : Fitted parameters ar
## either modifying the upper and lower limit boundaries, or else using
## multstart when fitting the model (see the function documentation).

## Warning in onetcm(t_tac = tacdata$Times/60, tac = tacdata$TAC, input = input, : Fitted parameters ar
## either modifying the upper and lower limit boundaries, or else using
## multstart when fitting the model (see the function documentation).

## Warning in onetcm(t_tac = tacdata$Times/60, tac = tacdata$TAC, input = input, : Fitted parameters ar
## either modifying the upper and lower limit boundaries, or else using
## multstart when fitting the model (see the function documentation).
```

```
## Warning in onetcm(t_tac = tacdata$Times/60, tac = tacdata$TAC, input = input, : Fitted parameters are
## either modifying the upper and lower limit boundaries, or else using
## multstart when fitting the model (see the function documentation).

## Warning in onetcm(t_tac = tacdata$Times/60, tac = tacdata$TAC, input = input, : Fitted parameters are
## either modifying the upper and lower limit boundaries, or else using
## multstart when fitting the model (see the function documentation).

## Warning in onetcm(t_tac = tacdata$Times/60, tac = tacdata$TAC, input = input, : Fitted parameters are
## either modifying the upper and lower limit boundaries, or else using
## multstart when fitting the model (see the function documentation).

## Warning in onetcm(t_tac = tacdata$Times/60, tac = tacdata$TAC, input = input, : Fitted parameters are
## either modifying the upper and lower limit boundaries, or else using
## multstart when fitting the model (see the function documentation).

## Warning in onetcm(t_tac = tacdata$Times/60, tac = tacdata$TAC, input = input, : Fitted parameters are
## either modifying the upper and lower limit boundaries, or else using
## multstart when fitting the model (see the function documentation).

## Warning in onetcm(t_tac = tacdata$Times/60, tac = tacdata$TAC, input = input, : Fitted parameters are
## either modifying the upper and lower limit boundaries, or else using
## multstart when fitting the model (see the function documentation).
```

```r
  # 2TCM
pbr28_long <- pbr28_long %>%
  mutate("2TCM" = pmap_dbl(list(tacdata, input, inpshift), fit_2tcm))
```

```
## Warning in twotcm(t_tac = tacdata$Times/60, tac = tacdata$TAC, input = input, : Fitted parameters are
## either modifying the upper and lower limit boundaries, or else using
## multstart when fitting the model (see the function documentation).

## Warning in twotcm(t_tac = tacdata$Times/60, tac = tacdata$TAC, input = input, : Fitted parameters are
## either modifying the upper and lower limit boundaries, or else using
## multstart when fitting the model (see the function documentation).

## Warning in twotcm(t_tac = tacdata$Times/60, tac = tacdata$TAC, input = input, : Fitted parameters are
## either modifying the upper and lower limit boundaries, or else using
## multstart when fitting the model (see the function documentation).

## Warning in twotcm(t_tac = tacdata$Times/60, tac = tacdata$TAC, input = input, : Fitted parameters are
## either modifying the upper and lower limit boundaries, or else using
## multstart when fitting the model (see the function documentation).

## Warning in twotcm(t_tac = tacdata$Times/60, tac = tacdata$TAC, input = input, : Fitted parameters are
## either modifying the upper and lower limit boundaries, or else using
## multstart when fitting the model (see the function documentation).

## Warning in twotcm(t_tac = tacdata$Times/60, tac = tacdata$TAC, input = input, : Fitted parameters are
## either modifying the upper and lower limit boundaries, or else using
## multstart when fitting the model (see the function documentation).

## Warning in twotcm(t_tac = tacdata$Times/60, tac = tacdata$TAC, input = input, : Fitted parameters are
## either modifying the upper and lower limit boundaries, or else using
## multstart when fitting the model (see the function documentation).
```

```
## Warning in twotcm(t_tac = tacdata$Times/60, tac = tacdata$TAC, input = input, : Fitted parameters are
## either modifying the upper and lower limit boundaries, or else using
## multstart when fitting the model (see the function documentation).

## Warning in twotcm(t_tac = tacdata$Times/60, tac = tacdata$TAC, input = input, : Fitted parameters are
## either modifying the upper and lower limit boundaries, or else using
## multstart when fitting the model (see the function documentation).

## Warning in twotcm(t_tac = tacdata$Times/60, tac = tacdata$TAC, input = input, : Fitted parameters are
## either modifying the upper and lower limit boundaries, or else using
## multstart when fitting the model (see the function documentation).

## Warning in twotcm(t_tac = tacdata$Times/60, tac = tacdata$TAC, input = input, : Fitted parameters are
## either modifying the upper and lower limit boundaries, or else using
## multstart when fitting the model (see the function documentation).

## Warning in twotcm(t_tac = tacdata$Times/60, tac = tacdata$TAC, input = input, : Fitted parameters are
## either modifying the upper and lower limit boundaries, or else using
## multstart when fitting the model (see the function documentation).

## Warning in twotcm(t_tac = tacdata$Times/60, tac = tacdata$TAC, input = input, : Fitted parameters are
## either modifying the upper and lower limit boundaries, or else using
## multstart when fitting the model (see the function documentation).

## Warning in twotcm(t_tac = tacdata$Times/60, tac = tacdata$TAC, input = input, : Fitted parameters are
## either modifying the upper and lower limit boundaries, or else using
## multstart when fitting the model (see the function documentation).

## Warning in twotcm(t_tac = tacdata$Times/60, tac = tacdata$TAC, input = input, : Fitted parameters are
## either modifying the upper and lower limit boundaries, or else using
## multstart when fitting the model (see the function documentation).

## Warning in twotcm(t_tac = tacdata$Times/60, tac = tacdata$TAC, input = input, : Fitted parameters are
## either modifying the upper and lower limit boundaries, or else using
## multstart when fitting the model (see the function documentation).

## Warning in twotcm(t_tac = tacdata$Times/60, tac = tacdata$TAC, input = input, : Fitted parameters are
## either modifying the upper and lower limit boundaries, or else using
## multstart when fitting the model (see the function documentation).

## Warning in twotcm(t_tac = tacdata$Times/60, tac = tacdata$TAC, input = input, : Fitted parameters are
## either modifying the upper and lower limit boundaries, or else using
## multstart when fitting the model (see the function documentation).

## Warning in twotcm(t_tac = tacdata$Times/60, tac = tacdata$TAC, input = input, : Fitted parameters are
## either modifying the upper and lower limit boundaries, or else using
## multstart when fitting the model (see the function documentation).

## Warning in twotcm(t_tac = tacdata$Times/60, tac = tacdata$TAC, input = input, : Fitted parameters are
## either modifying the upper and lower limit boundaries, or else using
## multstart when fitting the model (see the function documentation).

## Warning in twotcm(t_tac = tacdata$Times/60, tac = tacdata$TAC, input = input, : Fitted parameters are
```

```
## either modifying the upper and lower limit boundaries, or else using
## multstart when fitting the model (see the function documentation).

## Warning in twotcm(t_tac = tacdata$Times/60, tac = tacdata$TAC, input = input, : Fitted parameters are
## either modifying the upper and lower limit boundaries, or else using
## multstart when fitting the model (see the function documentation).

## Warning in twotcm(t_tac = tacdata$Times/60, tac = tacdata$TAC, input = input, : Fitted parameters are
## either modifying the upper and lower limit boundaries, or else using
## multstart when fitting the model (see the function documentation).
```

```
  # 2TCM1k
pbr28_long <- pbr28_long %>%
  mutate("2TCM1k" = pmap_dbl(list(tacdata, input, inpshift), fit_2tcm1k))
```

```
## Warning in twotcm1k(t_tac = tacdata$Times/60, tac = tacdata$TAC, input = input, : Fitted parameters a
## either modifying the upper and lower limit boundaries, or else using
## multstart when fitting the model (see the function documentation).

## Warning in twotcm1k(t_tac = tacdata$Times/60, tac = tacdata$TAC, input = input, : Fitted parameters a
## either modifying the upper and lower limit boundaries, or else using
## multstart when fitting the model (see the function documentation).

## Warning in twotcm1k(t_tac = tacdata$Times/60, tac = tacdata$TAC, input = input, : Fitted parameters a
## either modifying the upper and lower limit boundaries, or else using
## multstart when fitting the model (see the function documentation).

## Warning in twotcm1k(t_tac = tacdata$Times/60, tac = tacdata$TAC, input = input, : Fitted parameters a
## either modifying the upper and lower limit boundaries, or else using
## multstart when fitting the model (see the function documentation).

## Warning in twotcm1k(t_tac = tacdata$Times/60, tac = tacdata$TAC, input = input, : Fitted parameters a
## either modifying the upper and lower limit boundaries, or else using
## multstart when fitting the model (see the function documentation).

## Warning in twotcm1k(t_tac = tacdata$Times/60, tac = tacdata$TAC, input = input, : Fitted parameters a
## either modifying the upper and lower limit boundaries, or else using
## multstart when fitting the model (see the function documentation).

## Warning in twotcm1k(t_tac = tacdata$Times/60, tac = tacdata$TAC, input = input, : Fitted parameters a
## either modifying the upper and lower limit boundaries, or else using
## multstart when fitting the model (see the function documentation).

## Warning in twotcm1k(t_tac = tacdata$Times/60, tac = tacdata$TAC, input = input, : Fitted parameters a
## either modifying the upper and lower limit boundaries, or else using
## multstart when fitting the model (see the function documentation).

## Warning in twotcm1k(t_tac = tacdata$Times/60, tac = tacdata$TAC, input = input, : Fitted parameters a
## either modifying the upper and lower limit boundaries, or else using
## multstart when fitting the model (see the function documentation).

## Warning in twotcm1k(t_tac = tacdata$Times/60, tac = tacdata$TAC, input = input, : Fitted parameters a
## either modifying the upper and lower limit boundaries, or else using
## multstart when fitting the model (see the function documentation).
```

```
## Warning in twotcm1k(t_tac = tacdata$Times/60, tac = tacdata$TAC, input = input, : Fitted parameters a
## either modifying the upper and lower limit boundaries, or else using
## multstart when fitting the model (see the function documentation).

## Warning in twotcm1k(t_tac = tacdata$Times/60, tac = tacdata$TAC, input = input, : Fitted parameters a
## either modifying the upper and lower limit boundaries, or else using
## multstart when fitting the model (see the function documentation).

## Warning in twotcm1k(t_tac = tacdata$Times/60, tac = tacdata$TAC, input = input, : Fitted parameters a
## either modifying the upper and lower limit boundaries, or else using
## multstart when fitting the model (see the function documentation).

## Warning in twotcm1k(t_tac = tacdata$Times/60, tac = tacdata$TAC, input = input, : Fitted parameters a
## either modifying the upper and lower limit boundaries, or else using
## multstart when fitting the model (see the function documentation).

## Warning in twotcm1k(t_tac = tacdata$Times/60, tac = tacdata$TAC, input = input, : Fitted parameters a
## either modifying the upper and lower limit boundaries, or else using
## multstart when fitting the model (see the function documentation).

## Warning in twotcm1k(t_tac = tacdata$Times/60, tac = tacdata$TAC, input = input, : Fitted parameters a
## either modifying the upper and lower limit boundaries, or else using
## multstart when fitting the model (see the function documentation).

## Warning in twotcm1k(t_tac = tacdata$Times/60, tac = tacdata$TAC, input = input, : Fitted parameters a
## either modifying the upper and lower limit boundaries, or else using
## multstart when fitting the model (see the function documentation).

## Warning in twotcm1k(t_tac = tacdata$Times/60, tac = tacdata$TAC, input = input, : Fitted parameters a
## either modifying the upper and lower limit boundaries, or else using
## multstart when fitting the model (see the function documentation).

## Warning in twotcm1k(t_tac = tacdata$Times/60, tac = tacdata$TAC, input = input, : Fitted parameters a
## either modifying the upper and lower limit boundaries, or else using
## multstart when fitting the model (see the function documentation).

## Warning in twotcm1k(t_tac = tacdata$Times/60, tac = tacdata$TAC, input = input, : Fitted parameters a
## either modifying the upper and lower limit boundaries, or else using
## multstart when fitting the model (see the function documentation).

## Warning in twotcm1k(t_tac = tacdata$Times/60, tac = tacdata$TAC, input = input, : Fitted parameters a
## either modifying the upper and lower limit boundaries, or else using
## multstart when fitting the model (see the function documentation).
```

```r
  # Logan
pbr28_long <- pbr28_long %>%
  mutate("Logan" = pmap_dbl(list(tacdata, input, inpshift), fit_Logan))

  # mlLogan
pbr28_long <- pbr28_long %>%
  mutate("mlLogan" = pmap_dbl(list(tacdata, input, inpshift), fit_mlLogan))

  # MA1
pbr28_long <- pbr28_long %>%
```

```
  mutate("MA1" = pmap_dbl(list(tacdata, input, inpshift), fit_ma1))

  # MA2
pbr28_long <- pbr28_long %>%
  mutate("MA2" = pmap_dbl(list(tacdata, input, inpshift), fit_ma2))

pbr28_long <- pbr28_long %>%
  ungroup()
```

Now let's check the correlations between the estimated $BP_{ND}$ values.

```
pbr28_vtvals <- pbr28_long %>%
  ungroup() %>%
  select("1TCM", "2TCM", "2TCM1k", Logan, mlLogan, MA1, MA2)

art_corrplot <- ggcorrplot(cor(pbr28_vtvals), lab=T,
           colors = c("#6D9EC1", "white", "#E46726"),
           digits = 3, show.diag = F)

print(art_corrplot)
```

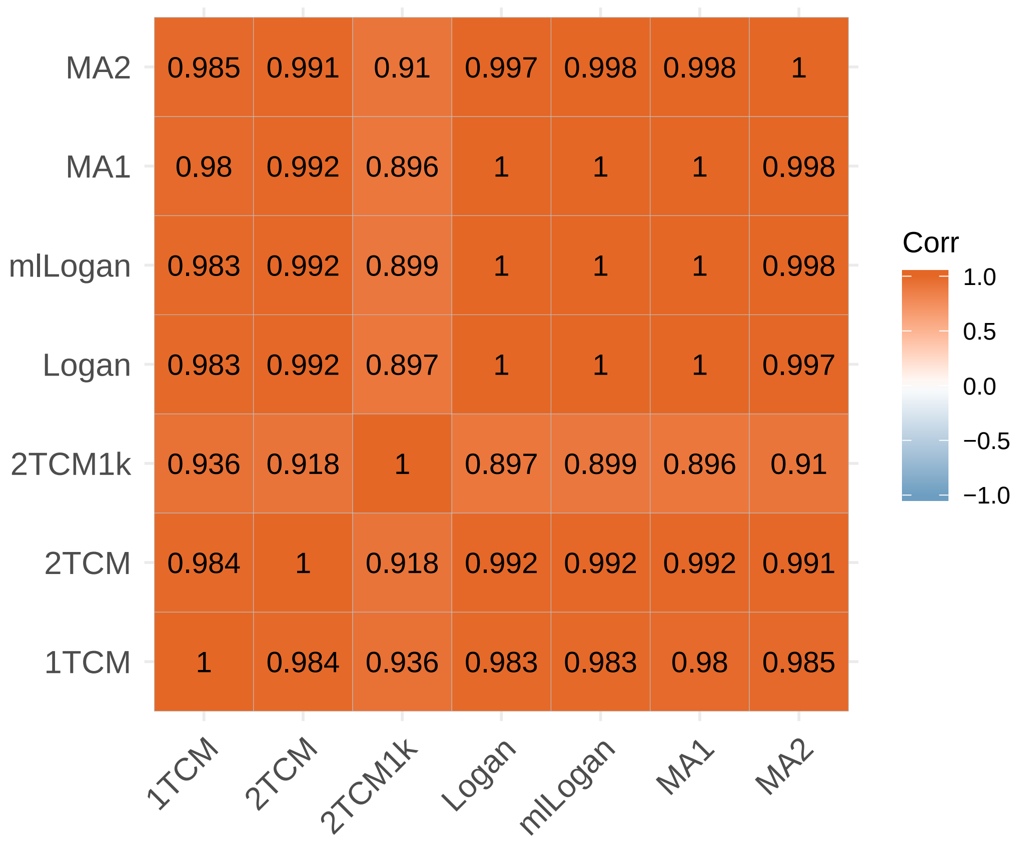
